# Domain segregated 3D chromatin structure and segmented DNA methylation in carcinogenesis

**DOI:** 10.1101/2020.01.13.903963

**Authors:** Yue Xue, Ying Yang, Hao Tian, Hui Quan, Sirui Liu, Ling Zhang, Yi Qin Gao

## Abstract

The three-dimensional (3D) chromatin structure, together with DNA methylation and other epigenetic marks, profoundly affects gene expression and displays abnormal behaviors in cancer cells. We elucidated the chromatin architecture remodeling in carcinogenesis from the perspective of spatial interactions between CGI forest and prairie domains, which are two types of megabase-sized domains defined by different sequence features but show distinct epigenetic and transcriptional patterns. DNA sequence strongly affects chromosome spatial interaction, DNA methylation and gene expression. Globally, forests and prairies show enhanced spatial segregation in cancer cells and such structural changes are accordant with the alteration of CGI interactions and domain boundary insulation, which could affect vital cancer-related properties. As the cancer progresses, a gradual increase of the DNA methylation difference between the two types of DNA domains is also observed for many different types of cancers. These observations are consistent with the change of transcriptional level differences of genes in these two domains, suggesting a highly-connected global structural, epigenetic and transcriptional activity changes in carcinogenesis.

## Introduction

Three-dimensional chromatin structure plays a vital role in gene regulation. The development of chromosome conformation capture (*1*) (3C) technology and its derived methods, such as Hi-C (*2*) and ChIA-PET (*3*), significantly improves our understanding of genome organization. For instance, the anchors of chromatin loops that frequently link enhancers and promoters are occupied by CCCTC-binding factor (CTCF) and cohesin complex in most cases (*4*). Such insulator structures can help maintain normal gene expression (*5, 6*). When it comes to cancer, many studies have revealed that mutations of CTCF binding sites and disruptions of insulated structures could result in dysregulation of gene expression (*6–8*), an intrinsic property in cancer. These studies mainly focus on local and specific genome regions. Besides, structural variants, such as deletions, inversions, translocations, are recurrent in multiple cancer types (*9*). Previous studies identified a positive correlation between translocation frequency and spatial proximity (*10*). A recent paper(*11*) has shown an integrative strategy to comprehensively detect these variants and captured numerous instances related to structural changes such as the fusion or loss of Topologically Associated Domains (TADs), the median size of which is several hundred kilobases. TADs were found to be largely conservative among different cell types (*12*). Nevertheless, unlike early embryonic development (*13*) and cell differentiation (*14*), the overall structural changes in carcinogenesis remain to be elucidated.

Along with aberrant 3D chromatin architecture, drastic genome-wide epigenetic changes also take place in carcinogenesis (*15, 16*), jointly influencing gene expression. Many studies have shed light on the stable epigenetic alterations associated with cancer cells, and DNA methylation was firstly and most widely studied (*17, 18*). There are mainly two types of general DNA methylation changes in cancer cells: global hypomethylation of late-replicating lamin-associated domains (LADs) (*19*) and hypermethylation of specific CpG islands (CGIs) (*20, 21*). Over ten thousands of publications reported DNA methylation changes as cancer biomarkers (*22*), but few have been applied to clinical diagnosis and treatment. While some studies provided examples to illustrate the regulation of between gene expression by DNA methylation, recently more evidences show that DNA methylation has little impact on gene expression but corresponds to chromosomal structural changes (*23, 24*). However, the correlation between changes of DNA methylation and cancer development and its relationship with chromosomal structural changes remain largely unknown. In principle, both chromosomal structure and epigenetic modifications can influence gene expression. Based on Hi-C contact map, the chromatin is divided into compartments A and B (*2*). Genes are enriched in compartment A and their expression levels are higher than those in compartment B. But there are many questions remain unanswered, e.g. what factors determine the compartment formation, what are the driving forces of compartment switch, and what are the roles of compartmentalization in cancer? Our previous study (*25*) showed that the compartment formation is strongly related to the genome composition. Based on the uneven distribution of CGIs, the whole genome was divided into two types of megabase-sized domains, CGI-rich domains (named as CGI forest domains) and CGI-poor domains (named as CGI prairie domains). These two types of domains, differing in sequence features, show distinct epigenetic and transcriptional patterns and overlap strongly with the compartments A and B, respectively. Furthermore, the cell-specific spatial contact and separation between these two types of domains are strongly coupled with various biological processes, such as early embryonic development (*26*), cell differentiation, and senescence (*27*). The main goal of this study is to understand the sequence dependence of various carcinogenesis marks and we found that forest and prairie behave significantly differently, including their distribution in compartments, CGI interactions, TAD formation, gene expression and epigenetic marks, especially DNA methylation, which is closely associated with development stage of cancer. The property difference between forest and prairie enlarges in cancer, consistent with their increased spatial separation in carcinogenesis. We also found that the regulation of gene expression depends on the sequence feature in a scale-dependent manner. In particular, a group of specific CGI genes that are originally marked by high repressive histone modifications become further hypermethylated, at the same time resulting in general repression.

## Materials and methods

### Source of methylome data

The whole-genome bisulfite sequencing (WGBS) data of methylomes were obtained from TCGA project and Gene Expression Omnibus, including 48 cancer samples and 17 matched adjacent samples. The reference genome is hg19. Normal liver and lung methylomes and those of their corresponding cancer cell lines were downloaded from Roadmap (*28*) and Encode Project (*29*) for combinatorial analysis of histone modifications and Hi-C contact. The description and references of the data sets are summarized in Supplementary Table S1. To ensure the credibility of the analysis results, in our calculation we only use CpG sites with coverage greater than three. DNA methylation level of each CpG site was given in percentage by

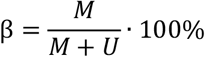

where M and U are the signal strength of methylated and unmethylated CpG, respectively. Besides, we calculated the normalized Z-score profile of histone modification signals within promoter and gene body for each gene,

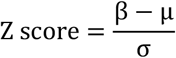

where μ and σ are the average and standard variation of signals on promoter or gene body of all coding genes, respectively. In this work, we focus on all protein coding genes which are downloaded from GENECODE release 19 (https://www.gencodegenes.org).

### Definition of CGI hypermethylation and hypomethylation

We calculate the methylation differences for 17 normal tissue samples and use 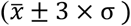 as the definition of CGI hypermethylation and hypomethylation, where 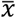 and σ refer to the average value and standard deviation, respectively. CGI hypermethylation or hypomethylation are defined as if the average methylation level of a CGI decreases or increases by a value of more than 0.3, respectively.

### F-P methylation difference (MDI)

Following our previous work (*25*), the methylation difference in open sea (regions beyond 4000 base pairs upstream and downstream of CGI) between neighboring forests and prairies is defined as:

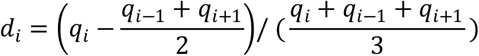

where *q_i_*, *q_i−1_*and *q_i+1_*are the methylation level for the *i*th domain and its two flanking domains.

### Definition of CGI genes

Promoter is defined as 1500 base pairs upstream of TSS (transcription start site) of a gene (*30*). When a gene or its promoter overlaps with CGI, it is defined as a CGI gene. The complement of CGI genes is nonCGI genes.

### Gene function analysis

Gene ontology (GO) enrichment analysis of all the given gene clusters in this work were conducted using the R package ClusterProfiler (*31*). Individual gene functions are obtained from GeneCards (https://www.genecards.org/). Immune related genes are obtained from AmiGO2 (http://amigo.geneontology.org/amigo) and oncogenes and tumor suppressor genes are downloaded from COSMIC (https://cancer.sanger.ac.uk/cosmic).

### Definition of tissue specificity for gene

The normalized RNA-seq data of GTEx project (*32*) was downloaded from https://zenodo.org/record/838734 (*33*). The tissue specificity of gene *i* in tissue *t* was defined as

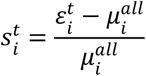

where 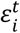 and 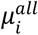 are the mean expression level of gene *i* in tissue t and all tissues examined, respectively. A gene with a tissue specificity value greater than 2 is defined as a tissue specific gene.

### Compartment identification

All human Hi-C data in this work were normalized by ICE method at a 40-kb resolution using the iced python package (*34*). Mouse cell cycle Hi-C data were normalized at 100-kb resolution and the reference genome is mm9. The identification of compartments A and B follows our previous work (*25*). To eliminate the influence of the centromere, the Hi-C matrix was disassembled into two parts, corresponding to p and q arms, and the eigenvalue decomposition was done within these two arms separately.

### Interaction strength

The 40-kb bin (in accordance with the resolution of Hi-C contact matrix) is identified as a CGI bin if it harbors at least one CGI, otherwise it is labeled as a nonCGI bin. With the above definition, each bin could spatially contact with four categories of DNA domains: CGI in CGI-rich domains (F-CGI), nonCGI in CGI-rich domains (F-nonCGI), CGI in CGI-poor domains CGI (P-CGI) and nonCGI in CGI-poor domains (P-nonCGI). The contact score between bin *k* and one of the four types of DNA segments *R_i_* is defined as

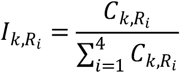

where *R_i_* is a vector consisting of the bins belonging to part *i, C*_*k,R*_*i*__ is the summation of all contact probabilities between bin *k* and *R_i_*. In this calculation, we deleted the self-contact elements.

### Contact probability and segregation factor as functions of genomic distance

The segregation factor was calculated as the ratio between contact probabilities of DNA domains of the same (F with F, or P with P) and different genome types (F with P), reflecting the extent of forest or prairie segregation. To identify contact loss in cancer cell line, we firstly calculated the average contact probability at the particular range of genomic distance for each bin for both cancer and its corresponding normal tissue. If this contact probability is higher than average level of all bins in normal cells but lower than average in cancer cells, then this bin is considered as contact loss.

### Definition of insulation score (IS)

For two neighboring regions *A_1_* and *A_2_*, the insulation score was defined as in Ref. (*35*),

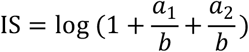

where *a_1_*, *a_2_* and *b* represent the mean contact probability inside *A*_1_ and *A*_2_ that between them, respectively. *A*_1_ and *A*_2_ can represent not only the forest and prairie domains, but also any two windows with the same size.

### Process of RNA-seq data

We downloaded counts formatted files from The Cancer Genome Atlas (TCGA) project for all available RNA-sequencing data of cancer and matched normal samples (summarized in Supplementary Table S1). Considering the property differences between normal and cancer cells, we calculated the relative expression levels for genes within each sample. The highest expressed gene possess highest relative expression level, namely, the amount of coding genes, and this value is equal in all samples. At the same times, the relative expression level is defined as zero for silenced genes.

## Results

### DNA methylation changes are closely related to cancer development

Consistent with previous studies, changes of DNA methylation from normal cells to cancer have two general characteristics: hypomethylation in the open sea (*30*) and hypermethylation in a subset of CGIs. Moreover, the extent of methylation changes appears to correlate with the stage of cancer development. At the early stage of carcinogenesis, little changes are observed in the methylation level for both CGI and open sea. As the cancer stage progresses, hypermethylation of CGIs occurs first, followed by hypomethylation of open seas. Similar trends of methylation changes are observed among a variety types of cancers, indicating the similarity in the development of different cancers or even potential common causes (Figure 1A and Supplementary Figure S1).

**Figure 1.**
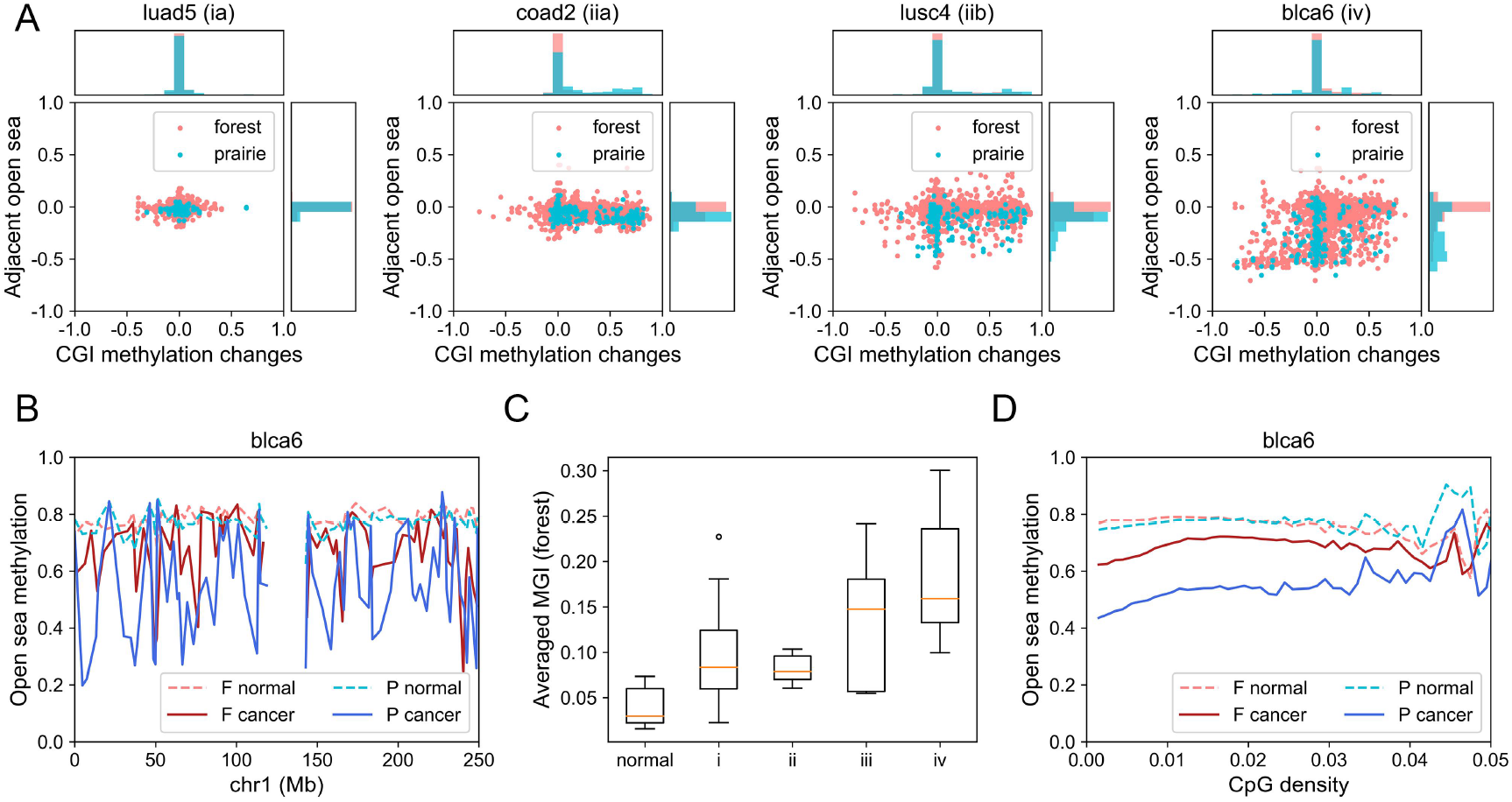
Methylation changes in carcinogenesis. (A) Scatter plots for changes of methylation level in CGIs and open seas. Each dot represents the methylation changes of a CGI (x axis) and its adjacent open sea (y axis) from adjacent normal samples to cancer samples on chromosome 1. The probability density of changes of CGI and open sea are shown on the top and right sides of the figure, respectively. (B) The average methylation level for CpGs located in open sea for every forest (F) or prairie (P) domain along the genomic sequence. (C) The averaged MGIs of all forest domains in normal samples and cancer samples in different stages. (D) CpG density and methylation level are calculated for all 1-kb beads in open sea. Each point on the curve shows the averaged methylation level for beads possess a given CpG density.

### Hypomethylation of open sea reflects the development of cancer

In most normal tissues, CpGs are mainly methylated in the open sea and the average open sea CpG methylation level in prairies (P) is slightly lower than that in forests (F). In carcinogenesis, open sea CpGs in prairies are more significantly hypomethylated than forests, leading to the increased methylation difference between forests and prairies (Figure 1B). The hypomethylation of the prairies gives rise to most of the PMDs observed earlier (*36*) (Supplementary Figure S2A). To quantify the difference between the open sea methylation levels of F and P domains, we calculated the averaged F–P methylation differences (MDI, see methods) for each sample and found that in normal tissues, the averaged MDIs for forests are always positive and that for prairies, negative, suggesting that the open sea methylation level of forests is in general higher than that of adjacent prairies. In cancer cells, averaged MDIs for forests become larger than their adjacent normal cells for almost every cancer sample (Supplementary Figure S2B). Remarkably, the averaged MDIs of forests generally increase with the aggravation of cancer, implying that the open sea methylation difference between forest and prairie domains can be used to reflect the stages of cancer (Figure 1C and Supplementary Figure S2C).

Furthermore, we found that the probability of hypomethylation increases with lowering CpG density in open seas of both forest and prairie domains (Figure 1D and Supplementary Figure S2D). In addition, the methylation level of prairie open seas is lower than that in forests even when they have the same CpG density. Such a result suggests that the prairie domains undergo more severe hypomethylation during carcinogenesis than the forest domains, suggesting that not only the local low CpG density, but also the surrounding sequence environment influences the methylation level of an open sea region.

### CGI hypermethylation and corresponding functional effects

In contrast to open sea CpG, the methylation level of the majority of CpGs in CGIs remain unchanged during carcinogenesis. Among those CGIs that do experience alternation of methylation, a larger proportion (6.1% of all CGIs on average) undergo hyper- than hypomethylation (2.2% of all CGIs) (Supplementary Table S2). Most of the hypomethylated CGIs are originally highly methylated in normal samples and they have typically low CpG densities and GC content, as well as small sizes, indicating that their similarity to open sea (Figure 2A, Supplementary Figure S3 and Supplementary Table S3). These results show that CpG density (at both CGI and forest/prairie length scales) is a major factor that determines the tendency for a CpG to become hypomethylated in carcinogenesis.

**Figure 2.**
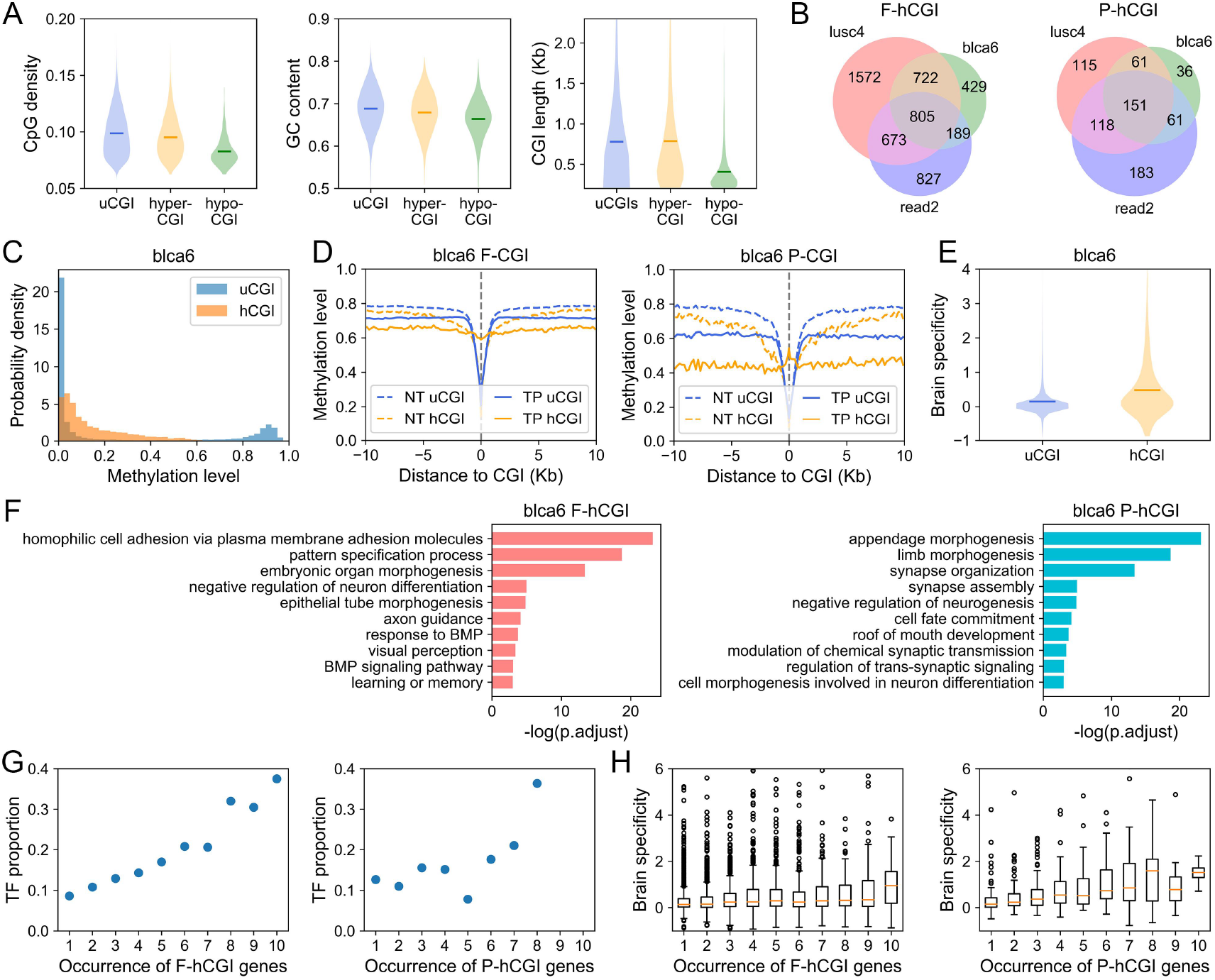
Changes of CGI methylation and corresponding functional characteristics. (A) CpG density (left), GC content (middle) and length (right) for methylation unchanged CGI (uCGI), hypermethylated CGI (hyper-CGI) and hypomethylated CGI (hypo-CGI), respectively. All with *P*-value < 10^−50^ by Welch’s unequal variance t-test between unchanged and hypomethylated CGIs. (B) The overlap of hypermethylated CGIs among three cancer samples. (C) The probability density of methylation level for uCGI and hCGI in blca6 (breast, normal) sample. (D) Methylation level of CGIs (at x=0) and their upstream (x<0) and downstream (x>0) CpGs. NT and TP refer to normal tissue and primary tumor, respectively. (E) The brain tissue specificity for uCGI and hCGI genes, *P*-value = 1.2 × 10^−30^ by Welch’s unequal variance t-test. (F) GO enrichment analysis for F-hCGI genes (left) and P-hCGI genes (right). (G) The proportion of transcription factors and (H) the brain specificity for hCGI genes which are observed in different accumulated numbers of samples.

Hypermethylation of specific CGIs is a common feature of all types of cancers examined, and is consistent with the previous studies (*37*). Moreover, these hypermethylated CGIs (hCGIs) are significantly conserved, which can be observed among both same and different types of cancers (Figure 2B). For instance, between urothelial bladder cancer and rectal adenocarcinoma, which are both at stage iv, hCGIs highly overlap. (*P*-value < 10^−322^ and < 10^−68^ by Fisher’s exact test for F-hCGI and P-hCGI, respectively). For most samples, the proportion of hypermethylated P-CGI is higher than F-CGI (10.6% of P-CGI, 5.7% of F-CGI on average, Supplementary Table S2). Notably, the original methylation level of hypermethylated CGIs in normal cells is in general higher than unchanged CGIs (uCGIs) except for those remain highly methylated in both normal and cancer samples (18.6% of uCGIs), indicating that hCGIs are already partially methylated in normal cells (Figure 2C and Supplementary Figure S3A). Moreover, open seas adjacent to hCGIs are more likely to be hypomethylated than those adjacent to uCGIs, suggesting a tendency of neighboring CGI and nonCGI becoming uniformly methylated in these regions (Figure 2D and Supplementary Figure S4A). This observation shows the existence of a long-distance correlation in DNA methylation along the genome, consistent with the earlier report (*38*).

In addition, genes of hypermethylated promoters or intragenic CGIs also show unique functional characteristics and histone modification patterns, which we will discuss in details later.

### Functions of CGI hypermethylated genes

Due to the conservation of hypermethylated CGI sites among different cancer types and stages, the functions of affected genes are also highly conserved. Taking sample blca6 as an example, the functions of forest hCGI genes are significantly enriched in cell adhesion, pattern specification, embryonic organ development and morphogenesis, including FOX, HOX, SOX, NKX, IRX genes (Figure 2F). Many of them are known to be transcription factors critical for embryonic development. These transcription factors are shared by a large number of cancer samples as the common sites of hypermethylation (Figure 2G). Except for developmental genes, the prairie hCGI genes are also noticeably enriched in brain and nerve functions, such as synapase organization, regulation of neurogenesis and chemical synaptic transmission (Figure 2F). As seen in Figure 2E and Supplementary Figure S4B, the brain tissue specificity for hCGI genes are higher than uCGI, for both forests and prairies, especially for hCGI genes common to different samples (Figure 2H). In addition, CGIs located in genes specific to other tissues could also be hypermethylated, but the brain related genes are most affected, probably partially because of the existence of the large amount of brain-specific CGI genes (Supplementary Figure S4C).

### Altered repressive marks for CGI genes in cancer cells

The hCGI genes were previously found to enrich H3K27me3 and repressed by polycomb complex in normal cells (*39*). We compare here the repressive histone mark signals in normal and cancer cells. Compared to uCGI genes, hCGI genes are not only enriched in H3K27me3, but also another repressive histone modification H3K9me3 in normal cells. Interestingly, the differences on these two repressive histone marks between hCGIs and uCGIs decrease or even disappear in cancer cell lines (Supplementary Figure S5 and S6). For instance, in the normal liver sample, H3K27me3 marks are significantly enriched in genes of hypermethylated CGIs in cancer. In the liver cancer cell line HepG2, H3K27me3 for promoters and gene bodies of forest uCGI genes becomes comparable to hCGI genes. In prairie, the histone marks on hCGI genes decrease significantly, leading to a diminishing difference between hCGI genes and uCGI genes. Similar phenomena are also observed from the comparison between lung and lung cancer cell line A549. The hCGI and uCGI genes also have similar H3K9me3 strengths in both lung and liver cancer cell lines, although these two types of CGIs have rather different levels of H3K9me3 in corresponding normal tissues. These results suggest that the overall epigenetic pattern of hCGI genes changes from repressive histone marks and low methylation to high methylation during cancer development, signaling the change of gene regulation mechanisms in carcinogenesis.

### Chromatin structure changes in cancer cell lines

#### Domain segregation becomes more pronounced in cancer cell lines

The chromatin structural differences between somatic and cancer samples are investigated in the following. We again use A549 cancerous lung cell line and Panc1 pancreatic cancer cell line as representative cancer samples, and compare them with somatic lung and pancreas samples. We also analyzed fresh leukemia and lung cancer samples, their chromatin structure characteristics are similar to cancer cell lines (data will be published somewhere else). Chromatin compartments in cancer and somatic samples differ in both sizes and extent of segregation. First, we used compartment vector components to divide the chromosome into compartments A and B (Supplementary Figure S7A). The size of compartment B increases in the two cancer cell lines when compared to their corresponding healthy tissues (Supplementary Table S5).

Compartment formation is seen to largely follow DNA sequence characteristics, namely forest and prairie (Figure 3A and Supplementary Figure S7C). DNA domains which belong to compartment A in both normal and cancer cell lines are composed primarily of forests (83% for A549 and 84%for Panc1) and common compartment B are mostly prairies (81% for A549 and 81% for Panc1), implying the intrinsic sequence preference for compartment formation. Moreover, changing from normal tissue to cancer cell lines, a portion of forests switch from compartment A to B and their CpG densities are lower than those of forests conserved in compartment A (Figure 3B and Supplementary Figure S7B). These observations indicate that in cancer cells compartment B, which constitutes mainly prairie domains, tend to “invade” forests with lower CpG densities. We then use the averaged compartment vector component 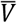 to quantify the DNA sequence preference of compartments (Supplementary Table S6). A high 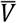 of a DNA domain implies that it has a high tendency to reside in compartment A. By comparing the 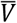 of CGI and nonCGI domains in forests and prairies, we found that in general, CGI domains and forests possess higher 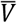 than nonCGI domains and prairies, respectively, for both normal and cancer samples. 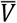s of sequences with different compositions all tend to decrease in carcinogenesis, suggesting an accumulation of compartment B in these samples. However, in carcinogenesis 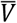 of CGI domains lowers more significantly than that of nonCGI domains, and CGI domains in prairies (P-CGI) shift the most significantly to compartment B among all four types of regions, demonstrating that there is a DNA sequence preference in the change of compartment segregation. The implication and biological function of such changes in chromatin compartmentalization will be analyzed as follows.

**Figure 3.**
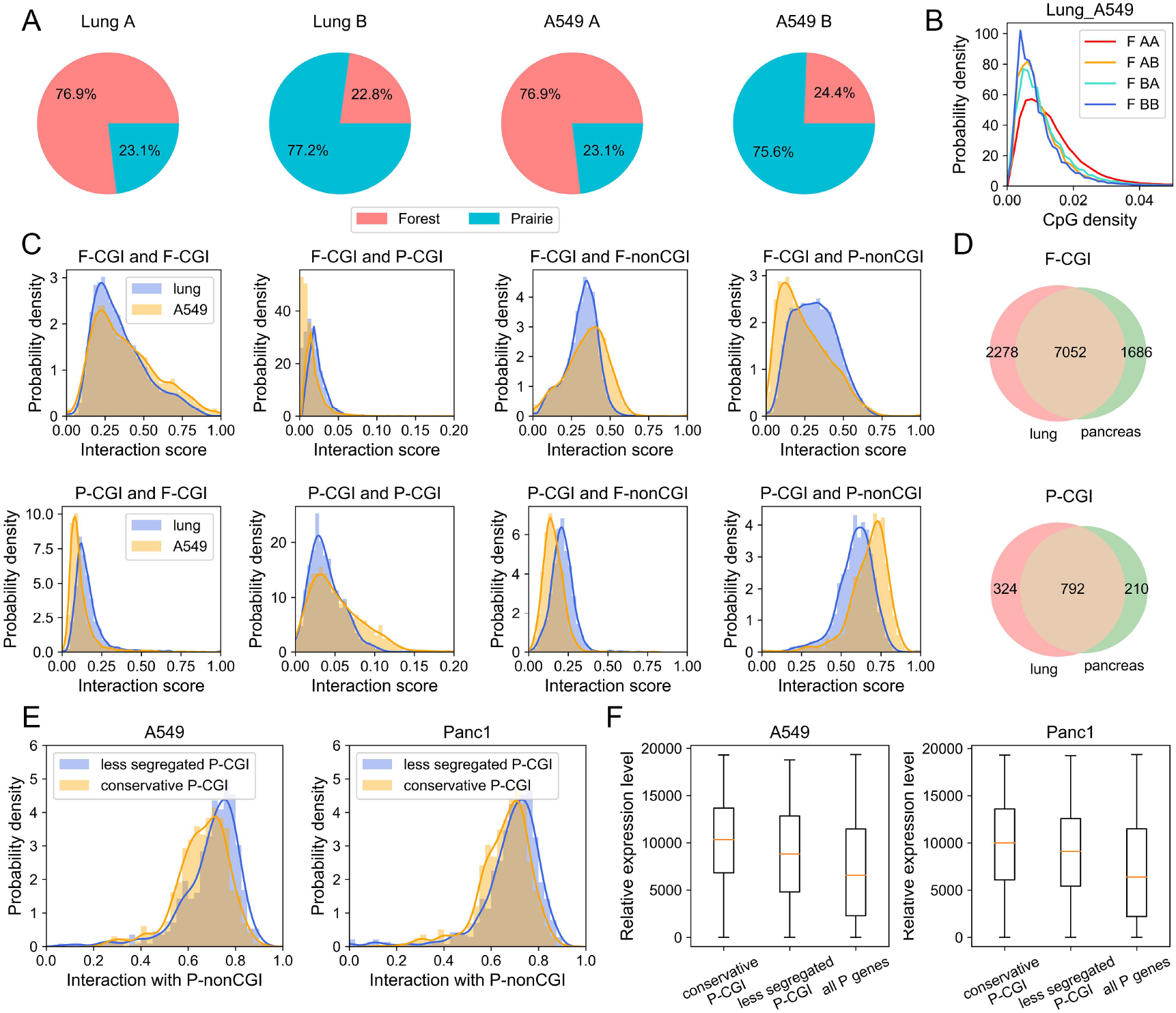
Compartmentalization and CGI aggregation in cancer cell lines. (A) The proportion of forest and prairie sequences in compartments A and B for lung and A549. (B) The probability density of CpG density for forest domains which belong to conservative compartment A (F AA) or B (F BB), as well as domains switch from A to B (F AB) or B to A (F BA) in carcinogenesis. (C) The interaction scores between F-CGI (top) or P-CGI (bottom) and the four types of domians (F-CGI, F-nonCGI, P-CGI, and P-nonCGI) in normal lung and A549. All *P*-values < 10^−90^ by t-test). (D) The overlap of aggregated CGIs between lung and pancreas. (E) The probability density of interactions with P-nonCGI for conservative P-CGIs and P-CGIs becoming less segregated in A549 (left) and Panc1 (right). *P*-value = *1.t* × 10^−7^ and *5.t* × 10^−5^ by Welch’s unequal variance t-test in A549 and Panc1, respectively. (F) The relative expression level for conservative P-CGI genes, less segregated P-CGI genes and all prairie genes in A549 and Panc1. Expression level for each gene is calculated by averaging expression levels over all LUAD samples and PAAD samples, respectively. *P*-value = *1.7* × 10^−5^ and 0.022 between conservative and decrease P-CGI genes by Welch’s unequal variance t-test in lung and pancreas, respectively.

#### CGI aggregation in carcinogenesis

We next focus on CGIs and investigate its 3D contact with other genomic components in both normal and cancer cells and try to reveal the underlying biological implications. We firstly calculated the interaction scores (see methods) for both normal cells and cancer cell lines. Taking lung as an example, from normal to cancer, contacts between the same genome types (F-CGI and F-CGI, F-CGI and F-nonCGI, P-CGI and P-CGI, P-CGI and P-nonCGI) all increase, accompanied by the reduced contacts between different genome types (F-CGI and P-CGI, F-CGI and P-nonCGI, P-CGI and F-CGI, P-CGI and F-nonCGI) (Figure 3C). These results clearly show that the two types of sequences, forests and prairies, become more segregated in the 3D space, as one changes from normal to cancer cells. NonCGI DNA regions also display a similar tendency (Supplementary Figure S7E).

Similar results are also obtained for pancreas cancer (Supplementary Figure S7D, S7F). Intriguingly, the F-CGIs and P-CGIs having strong contact with the same types (between F-CGIs and between P-CGIs) in cancer are highly conserved between lung and pancreas (*P*-value < 10^−300^ and – 10^−20^ by Fisher’s exact test in forest and prairie, respectively, Figure 3D). For the convenience of discussion, we hereinafter name these common CGIs conservative CGIs. Notably, in cancer, compared with less segregated P-CGIs, the conservative P-CGIs show significantly lower contact probability with P-nonCGI regions (Figure 3E), the less active chromatin domains. This result indicates that the aggregation of P-CGI during carcinogenesis may result in a more open and active environment (although within compartment B) which attributes to the change of gene expression level (Figure 3F).

To link the structural changes mentioned above to biological functions, we divided genes containing conservative CGIs into two groups based on whether their expression levels increase or decrease in carcinogenesis. We found that up-regulated forest genes in lung cancer cells are associated with chromosome segregation and glycosylation. The former is relevant to cell division and the latter, a modification that adds glycan to proteins or other molecules and could further affect cell communications and interactions, is known to play vital roles in cancer development and progression (*40*). In pancreas, besides the functions mentioned above, up-regulated genes are also of functions such as embryonic organ development and morphogenesis, suggesting a strong relationship between carcinogenesis and early embryo development (*41*), which is worthy of further investigations. In the conservative prairie regions, functions of up-regulated genes in lung cancer cells are related to Wnt signaling pathway and cell division. Wnt signaling, a group of signal transduction pathways, has been linked to cancer owing to its role in regulating development (*42*). Such genes found in the analyses on pancreas samples act on epithelial to mesenchymal transition (EMT) and organ development, contributing to the cell growth and invasiveness in carcinogenesis(*43*). These analyses thus suggest that the spatial aggregation of CGIs and functional changes are highly correlated in tumorigenesis (Supplementary Table S7 to S10).

#### General architecture of chromatin in cancer cell

The structure analyses above indicate that significant chromatin structure changes occur at both CGI and compartment scales in carcinogenesis. Next, we try to investigate the chromatin structural changes at a wide range of scales. We first analyzed the contact probability change at varied genomic distances (see methods), and observed that overall the contact probability decays faster as a function of genomic distance for cancer cells than normal cells, which indicates a loss of long-range spatial contacts in carcinogenesis (Supplementary Figure S8A). Further investigation revealed that the F-P contact is lower than F-F and P-P contacts at nearly all sequential distances, indicating the overall separation between these CGI-rich and CGI-poor domains (Figure 4A and Supplementary Figure S8B). The contact probability calculated for the normal lung cell decays following almost a single power-law in the genomic distance range of hundreds of kilo- to several mega-bases (slope = −0.76 and −0.69 for F-F and P-P contacts, respectively), indicating a relatively uniform contact probability scaling property for normal tissues. In contrast, the cancer samples exhibit a scale separation in contact probability decay curve, with a slower decay for both F-F (slope=−0.56) and P-P (slope=−0.55) contacts than corresponding somatic samples at distances shorter than 400kb (F-F contact) and 800kb (P-P contact), and a steeper decay (slope=−1.37 and −1.26 for F-F and P-P contacts, respectively) at large distances (Supplementary Figure S8C and S8D).

**Figure 4.**
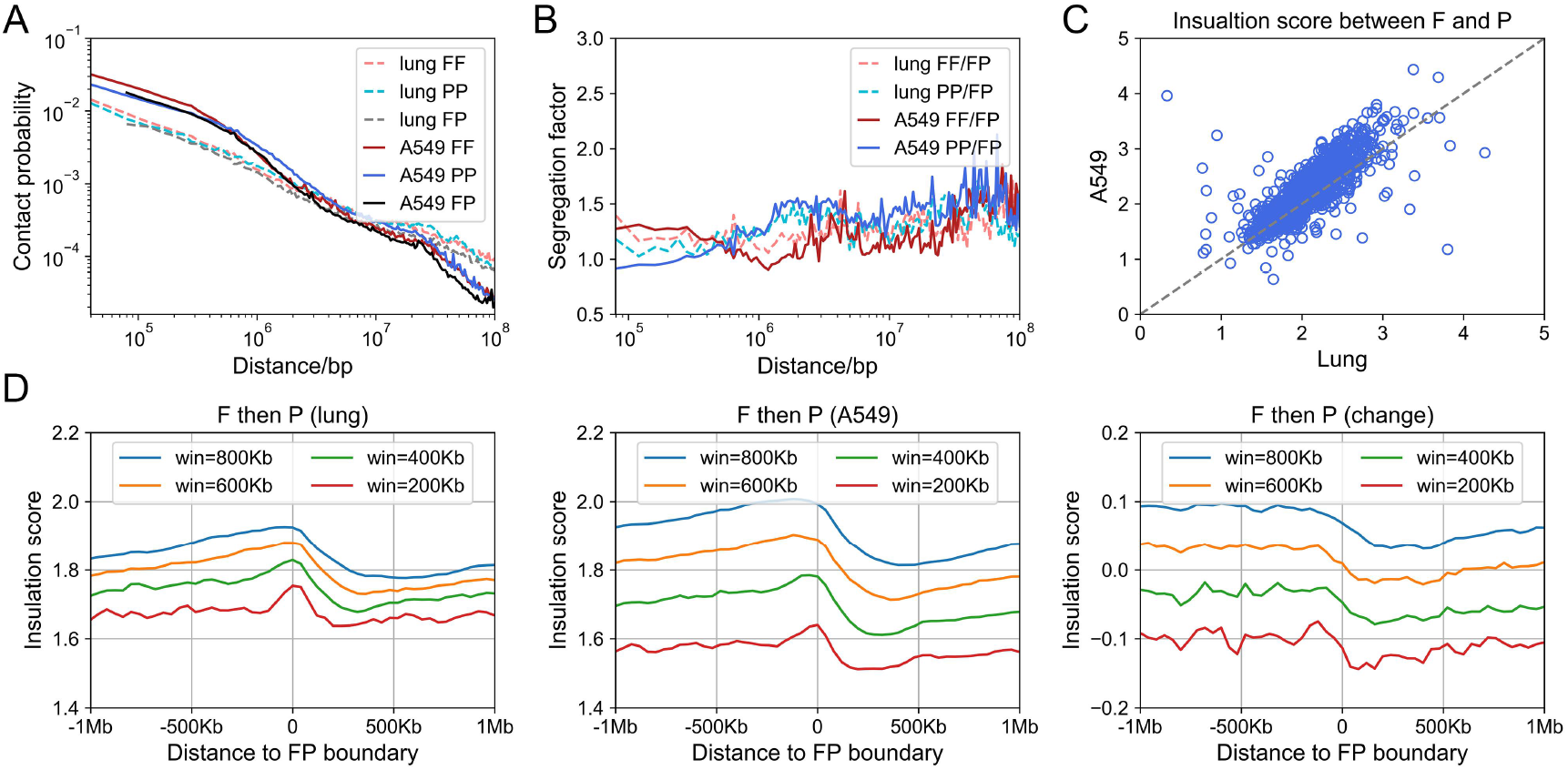
General chromatin architecture in cancer cell lines. (A) The contact probability between forests and forests (FF), prairies and prairies (PP), forests and prairies (FP) at varied genomic distances for lung and A549 (chromosome 1 is used as an example). (B) The segregation factor at varied genomic distances for lung and A549. (C) The insulation score between adjacent forest and prairie domains in lung and A549. (D) The insulation scores for 40-kb beads around F-P boundary at different window sizes in lung (left), A549 (middle) and the difference between A549 and lung (right). The data are aligned so that the forest domain is left to the boundary (value 0).

To further compare the relative contact strength of forests and prairies at varied distances, we defined and calculated the segregation factor (see methods). A high segregation factor for a DNA segment (e.g., of 40-kb) indicates that it prefers to contact with domains of the same type over those of a different type at that given genomic distance. For both normal and cancer samples, the segregation factor is almost always greater than 1 at all genomic distances, suggesting an overall F-P domain separation (Figure 4B). At short distances (less than 500kb) the segregation factor is higher for forests than for prairies. As the genomic distance increases, its value decreases for forests and increases for prairies, changing from normal to cancer cells. These observations indicate that forest domains have strong contacts at short distances, especially those within the same forest. Spatial contacts between F-domains are weak at a genomic distance of ~1Mb. In contrast, contacts between nearby prairie DNAs are weak but when the genomic distance increases to millions of kilobases, prairie domains tend to interact frequently.

Intriguingly, a number of forest genes lose contact with other forest domains at genomic distances ranging from 600K to 2M in cancer cells, contributing to the weakened segregation factor for F-domains at ~1M. These genes are heavily shared between A549 and Panc1 (*P*-value < 10^−300^ by Fisher’s exact test) and many are related to the immune process (Supplementary Figure S8E, S8F). For instance, 30% of the genes related to antigen processing and presentation and 26.2% of immune system genes are involved in the F-F contact loss in A549 cancer cell line. In Panc1 cell line, the proportions are 28.0% and 26.5% (Supplementary Table S11 and S12). For example, a forest gene RELA, which is a proto-oncogene and subunit of NF-κB, is found to lose contact with forest domains in both tumor samples. On the one hand, dysregulation of NF-κB is a hallmark of cancer. On the other hand, it can promote genetic and epigenetic alterations, change cellular metabolism, directly and indirectly control inflammation, cancer cell proliferation and survival, epithelial-to-mesenchymal transition (EMT), invasion, angiogenesis and metastasis (*44*). Common genes also include kinesins, the misregulation of which are involved in cancer pathogenesis, such as uncontrolled cell growth and metastasis (*45, 46*). At the same time, a group of growth factors are also involved in this chromosome structure changes and the relationship between them and how they contribute to cancer initiation and development remains to be further investigated.

We also used the insulation score (IS, see methods) to explore the structure changes in carcinogenesis. From the perspective of the domain level, the IS between adjacent forests and prairies was significantly larger in tumor than in normal cell (*P*-value = 4.12 × 10^−83^ and 6.28 × 10^−10^ by t-test for lung and pancreas, respectively, Figure 4C and Supplementary Figure S8H), again hinting the formation of a structure with F and P-domains significantly separated. We next investigated the spatial insulation around F, P-domain boundaries at varied window sizes (Figure 4D and Supplementary Figure S9). For both normal and cancer cell lines, insulation scores of bins located in forests are generally higher than those in prairies, indicating more local interactions within forest, accordant with our previous work that forests are mainly composed of type A whereas prairies are type B (*25*) Furthermore, the IS in both forest and prairie in tumor is smaller than that in normal cells at small window sizes (e.g., 200kb), indicating a more homogeneous distribution of contact around the main diagonal of Hi-C matrix. As the window size increases, forests and prairies display distinctly different insulation behaviors. For forests, the cancer IS values become larger than the corresponding values in normal tissues when the window size is larger than ~500kb, indicating that the interactions in forest domains becomes dominated by local contacts. In contrast, the IS values in prairie domains are smaller in cancer samples compared with normal cells at larger range of window sizes than that in forest, corresponding to strengthened contacts between linearly distant genomic regions within prairie domains.

The differences between cancer cell lines and normal cells in terms of DNA contacts, segregation factors and insulation scores provide a consistent picture for chromatin structural change in carcinogenesis. In cancer samples, the normalized short-range contact increases. The segregation factor difference between forests and prairies also increases at short genomic distances, indicating an increased difference between forest and prairie local 3-D structures in carcinogenesis. Changing from normal to cancer cells, the segregation factor of forest decreases more sharply in short distances than that of prairies, indicating contacts in forests to become more local, consistent with the change of insulation score. In contrast, the segregation factor of prairies increases more significantly with the genomic distance, showing a prominent peak at ~800kb, indicating the loss of local contacts in the expense of long range intra- and inter-domain prairie contacts. Interestingly, the change of contacts between F-domains and those between P-domains occur at genomic distances corresponding to their TAD sizes, respectively, indicating the improved formation of TADs and reduced inter-TAD contacts in both forests and prairies. As the size of compartment B increases in cancer and forests seldomly occupy this compartment, the higher segregation factor for prairie and lower factor for forest at long distances indicate the spatial clustering of distant prairies but not forests.

### Relationship between DNA methylation and chromosomal structure

#### CpG density dependence for DNA accessibility and methylation

It’s well known that the unmethylated CGI is in general free of nucleosomes and more accessible to the transcription factors compared to methylated CGI and other genomic regions. In an earlier study, we showed that DNA methylation of the open sea is correlated to the chromatin 3D structure. All these results suggest the importance of CpG density on CpG methylation and the openness of chromatin. Therefore, we divide DNA into four groups (Groups I, II, III, IV) according to their CpG density per thousand base pairs ((2.0%, 20.1%], (1.0%, 2.0%], (0.5%, 1.0%] and (0, 0.5%], respectively). (Beads located in CGIs mostly belongs to the first group as the minimum of their CpG density is 2.4%.) We then analyzed DNase I hypersensitivity and corresponding methylation data for liver and lung cancer cell lines (HepG2 and A549, respectively), as well as for all available somatic normal tissues.

In normal cells, DNase I hypersensitivity of group I is slightly lower than other groups. In general, for normal samples DNase I hypersensitivity decreases slowly with the increase of CpG methylation level regardless of CpG density. However, in cancer cells, regions with high CpG densities and low methylation levels are much more accessible than other regions, and the DNase I hypersensitivity decreases to nearly 0 when the CpG density is lower than 0.02 or the methylation level is higher than 0.5 (Figure 5A and Supplementary Figure S10).

**Figure 5.**
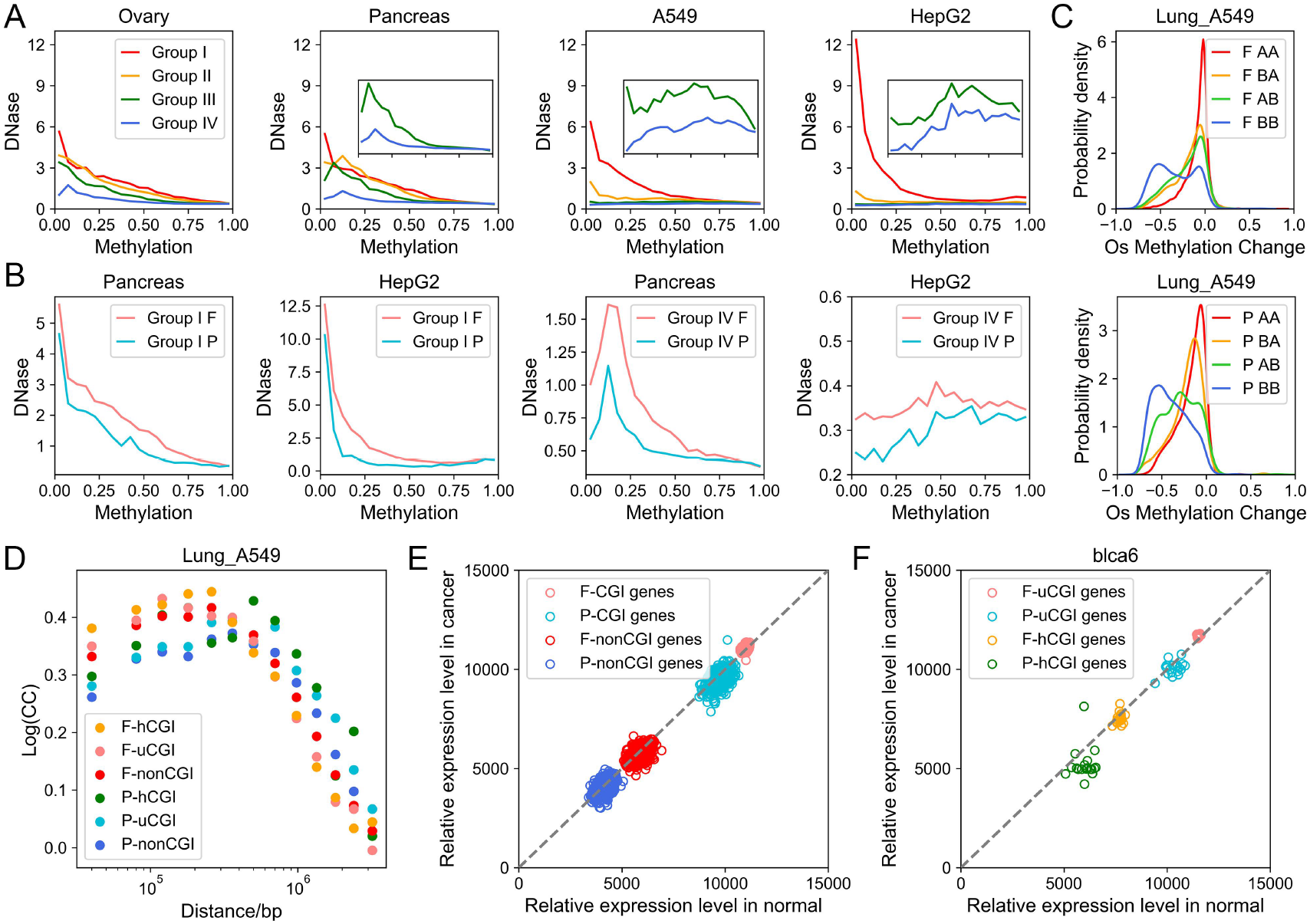
Association between methylation and chromosome structure and gene regulation. (A) Average DNase signal at various methylation levels. CpG density and CpG methylation level are calculated at a 1-kb resolution. Bins are divided to groups I, II, III or IV according to their CpG density (2.0%, 20.1%], (1.0%, 2.0%], (0.5%, 1.0%] or (0, 0.5%], respectively. Beads located on forest or prairie domains are shown separately in (B). (C) The probability density of open sea methylation changes for domains in forests (top) and prairies (bottom). (D) Contact change (CC) from lung to A549 at various genomic distances. (E) Average relative expression level for 675 pairs of normal-cancer samples. Each point represents the average expression level of genes belonging to a gene category in one sample. (F) Difference of average relative expression level for hCGI and uCGI in 19 pairs of BLCA samples.

Notably, with the decrease of CpG density, the relationship between DNase I hypersensitivity and methylation gradually switches from a negative to a positive correlation in tumor cells, indicating that genomic regions of very low CpG densities but high methylation are largely accessible. We also found that DNase I hypersensitivity in forests is constantly higher than that in prairies for any given CpG density and methylation level, and in both normal and cancer cells, consistent with the forest being in a more open and active environment (*25*). Remarkably, the DNase I hypersensitivity in prairies decreases more quickly than that in forests with the decrease of CpG density in cancer cells. Therefore, a positive correlation between DNase I hypersensitivity and methylation level persists in a larger DNA density range in prairies than that in forests (Figure 5B and Supplementary Figure S10).

#### Methylation of open sea is associated with chromatin structure

To obtain more details on the relationship between chromosomal structure and open sea methylation, we divide the genome into four groups: regions switch from compartment A to compartment B in tumorigenesis (AB), regions switch from compartment B to compartment A (BA), and those remain as A (AA) or B (BB) in both normal and tumor cells. The methylation level of AA regions remain largely unchanged and BB regions undergo the strongest demethylation, indicating that the genomic silent regions are more likely to be demethylated. Interestingly, AB regions are demethylated to a larger extent than BA, indicating that open sea demethylation tends to occur in the repressed domains of cancer cells rather than those of normal cells (Figure 5C).

There are several possible reasons behind hypomethylation and enlarged difference between forest and prairie open sea methylation. It was reported that CpG loci with multiple CpG sites in the surroundings are more efficiency methylated by DNMT1 (*47*), indicating that the local sequence feature partly contribute to the hypomethylation. However, the sequence property in the large scale (forest or prairie) is also likely to affect DNA methylation. In fact, prairies tent to undergo more drastic hypomethylation than forest regions even when they have the same local CpG density (Figure 1D). We also examined the sequence environment effects on solo-WCGW (‘solo’ refers to the CpGs with no neighboring CpGs and ‘W’ indicates A or T nucleotide), which is reported to be the most hypomethylation-prone sites in carcinogenesis (*24*) (Supplementary Figure S11). Notably, solo-WCGWs located in prairies also have a lower methylation level in normal cells and undergo more drastic demethylation in carcinogenesis compared to those in forests, further illustrating the importance of the sequence environment. A possible explanation for these observations is that cancer cells undergo more frequent cell cycles than normal cells, resulting in insufficient methylation and thus a global hypomethylation. Due to the domain segregation and resulted different accessibility of forests and prairies, this hypomethylation is more likely to occur in the latter domains, enlarging the methylation difference between forests and prairies. Such a mechanism is also consistent with previous findings of hypomethylation in aging cells, and the extent of hypomethylation being proportional to the replication timing of the regions and the cell division rate of a tissue (*24, 48*). In turn, the large methylation difference is expected to lead to a higher degree of segregation between forests and prairies.

Therefore, in general, methylation of open seas appears to reflect the chromosomal structure (whereas since CGIs are crucial for gene regulation, their methylation level is more directly related to the expression of genes). To further understand the relationship between CpG methylation and chromatin structure, we next investigate how the 3-D structure (in terms of Hi-C contacts) changes for hypermethylated CGIs in carcinogenesis.

#### CGI hypermethylation is associated with distinct chromatin structure between forest and prairie

We analyze the insulation score (see methods) of hypermethylated CGIs and unchanged CGIs of normal lung sample and A549. In all these samples, compared to F-uCGIs, F-hCGIs tend to contact with F-hCGI and F-uCGI rather than nonCGI regions, including both F-nonCGI and P-nonCGI. Changing from normal cells to cancer cell lines, both F-uCGI and F-hCGI experience an increase of contacts with forests and lowered contacts with prairies. At the same time, both P-uCGI and P-hCGI are more likely to form more contacts with prairie than with forest domains (Supplementary Figure S12A, S12B). These results are consistent with the cancer-related spatial segregation between forests and prairies. Furthermore, in normal cells, P-hCGIs and P-uCGIs tend to contact with domains of similar properties (P-hCGIs and P-uCGIs), respectively. To be more quantitative, we calculated the contact ratio between uCGI and hCGI within the same type of domains. For instance,

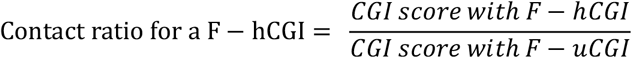

From normal cells to tumor cell lines, the contact ratios for F-hCGIs, F-uCGIs, P-hCGIs and P-uCGIs tend to increase (Supplementary Figure S12C). Therefore, CGI aggregation not only occurs between those of the same sequence environment (forests or prairies), but also between CGIs with similar methylation states in carcinogenesis.

To obtain a more complete picture of genomic contact formation, we next calculate the contact change (*CC*) between normal lung and A549 samples:

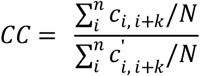

Where n refers to all the bins of the concerned sequence type and N refers to the number of these bins. At the given genomic distance k, *c_i, i+k_* is the contact probability between loci i and i+k in the Hi-C map of normal lung and 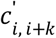, A549.

As seen from Figure 5D, *CC* is higher for F-hCGIs than for F-uCGIs at short linear genomic distances. At long distances, it is higher for P-hCGIs than for P-uCGIs. These results indicate that compared to normal cells, F-hCGIs in tumor cells are more likely to form local contacts and P-hCGIs are more likely to form distant interactions along the linear genome.

### Regulation of gene expression in carcinogenesis

Next, we examine whether the change of gene expression in carcinogenesis also shows a DNA sequence dependence. We first obtained 675 pairs of transcriptome (cancerous versus adjacent normal tissues) from TCGA (http://cancergenome.nih.gov), and compared their averaged transcription levels in CGI and nonCGI regions and in forests and prairies (see methods) (Figure 5E). CGI genes (genes with CGIs on their promoter or body, see methods) are on average ranked more highly in expression than nonCGI genes (*P*-value all < 10^−300^ for F and P genes in normal and cancer samples by Welch’s unequal variance t-test), and the averaged expression levels of forest genes are constantly higher than prairie ones for both CGI and nonCGI genes (*P*-value all < 10^−300^ for CGI and nonCGI genes in normal and cancer samples by Welch’s unequal variance t-test). These results show that the sequence property of not only the different components of the genes but also their surrounding sequences (especially, whether they reside in forest or prairie domains) could have a significant influence on their expression. Compared to their corresponding adjacent tissues, the averaged expression levels of cancer samples are higher for CGI genes in forests (*P*-value = 9.75 × 10^−65^ by t-test), and lower for those in prairies (*P*-value = 2.87 × 10^−67^ by t-test), showing a cancer-related increased difference between forests and prairies at the expression level. Notably, the change of gene expression and spatial segregation is only significant for CGI genes, whereas for nonCGI genes this difference between cancer and normal samples is not observed, consistent with the similar 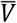 values seen in nonCGI regions of cancer and normal cells.

#### CGI hypermethylation globally repress gene expression

CGI hypermethylation has been argued to have a strong effect on gene expression.(*49*) Firstly, we explore the average effect of CGI hypermethylation and analyzed the relative expression level of 19 bladder urothelial carcinoma (BLCA) patients and 48 lung squamous cell carcinoma (LUSC) patients. The uCGI and hCGI genes are defined by the methylation changes of sample blca6 and lusc4, respectively. The averaged expression level of CGI genes that harbors hCGI are found to be significantly lower than genes with unchanged CGI methylation (*P*-value = 6.72 × 10^−33^ and 4.30 × 10^−29^ for F-hCGI and P-hCGI genes in BLCA normal samples by Welch’s unequal variance t-test, respectively. *P*-value = 1.01 × 10^−73^ and 1.68 × 10^−80^ in LUSC normal samples). The expression level of these genes further decreased in malignant cells (*P*-value = 8.72 × 10^−3^ and 9.34 × 10^−4^ for F-hCGI and P-hCGI genes in BLCA by t-test, respectively, *P*-value = 1.09 × 10^−7^ and 4.55 × 10^−15^ in LUSC), consistent with the general role of CGI hypermethylation in repressing expression (Figure 5F and Supplementary Figure S13A). Next, we investigate the influence of methylation on individual genes and analyze the relative expression level of four normal-cancer TCGA samples the WGBS data of which are also available. Surprisingly, we found not all CGI hypermethylated genes are repressed in carcinogenesis and a number of genes are even significantly upregulated. At the same time, the hypermethylation and expression level changes of cancer related oncogenes and repressors appear to be uncorrelated, indicating that the regulation of gene expression, as well as oncogenes and repressor genes, may involve factors more than just the methylation reprogramming in carcinogenesis (Supplementary Figure S13B). On the other hand, the average expression level of F-hCGI genes and P-hCGI genes decrease in carcinogenesis and the number of down-regulated genes are higher than up-regulated for most samples (except for luad5, which has very few hCGIs), consistent with the general repression effect of methylation (Supplementary Table S13).

In conclusion, gene expression appears to be regulated by the sequence at both forest/prairie (Mb) and CGI/nonCGI (kb) levels. In addition, CGI hypermethylation in cancer executes an overall repression on genes the expression levels of which are relatively low in normal cells, further enlarging the difference between the highly and lowly expressed genes. However, down-regulation does not occur to all such genes, indicating a complicated relationship between DNA methylation and individual gene expression.

## Conclusion and discussion

Genetic alterations at highly varied frequencies and complex combinations make cancer treatment difficult. In spite of the high heterogeneity, cancer cells consistently exhibit a number of similar characteristics, such as uncontrolled growth and proliferation. The importance of chromosomal structure and epigenetic modifications on cell function has been widely discussed and a large number of regulatory elements have been discovered, but the causal link between them remain unclear. In the present study, we focused on the epigenetic similarities between different types of cancer and performed an integrated analysis of chromatin DNA methylation, 3-D structure, DNase hypersensitivity and gene expression in carcinogenesis. In particular, to make connections to genetics, we try to identify the sequence dependence in these epigenetic changes. At a “local” scale, CGI and nonCGI regions present different DNA CpG methylation properties as well as transcriptional activities. Based on the uneven distribution of CGI, the genome can be divided into megabase-sized domains, CGI-rich domains (forests) and CGI-poor domains (prairies). The former possesses higher chromatin openness, gene density, transcription activity and open sea methylation level than prairies. Moreover, sequence provides a basic framework for the general phase separation that euchromatins are mainly formed by forests and heterochromatins are mainly formed by prairie, while variation among tissues is related to tissue specificity. In this work, we further illustrated that the discrepancy between forests and prairies is more than sequence difference. The two types of DNA domains represent two distinct genetic environments and the functional differences between them further increase in cancer cells.

For the overall chromosomal structure, compartment formation is strongly influenced by DNA sequence and potentially repressive regions tend to accumulate in compartment B. The sequential preference in the re-establishment of compartments and increased homotypic contacts indicates enhanced segregation of the two types of linear sequences in cancer. From normal to cancer cell lines, several notable chromatin structural changes can be observed: (1) The chromatin openness difference between high CpG density/unmethylated sites and low CpG density/methylated sites become more prominent. Although in normal cells, the DNase I hypersensitivity of forest is always higher than prairie even for regions with the same CpG density and methylation level, this difference between forests and prairies becomes more prominent in cancer cell lines. (2) Forests become more clustered at short distances and contacts within prairies tend to become more dispersed. (3) The isolation between forests and prairies increases. In particular, the contact probability between F-CGI and P-CGI significantly decreases in cancer samples compared to normal cells. The methylation levels of forest and prairie open seas of the same CpG density are also different. In carcinogenesis, prairies tend to undergo more drastic hypomethylation, which largely reflect the chromosome structural characteristics discussed above. The HiC data shows that the change of chromatin structure is likely driven by the aggregation of prairies, consistent with the finding that attractions between heterochromatic regions are crucial for the formation of compartment (*50*), and facilitated by the large number of cell cycles the cells experienced.

From the perspective of CGI, during carcinogenesis, CGIs within the same genome type (forest or prairie) tend to aggregate and this aggregation is highly conserved between lung and pancreas. Such structural changes are found to correlate with functions corresponding to carcinogenesis and cancer development. However, the mechanisms of these changes are not clear and the gene regulatory networks in cancer need to be further investigated. We speculate that since many transcription factors bind CGIs rich regions, the higher spatial contacts within CGIs may provide an open and active environment for related genes’ transcription in cancer (e.g., through a liquid-liquid phase separation mechanism(*51*)). The methylation state of CGI in genes also is well-known to influence gene expression, although we found that some hypermethylated genes become actually activated. The influence of CGI methylation could also have a 3-D structural component: it was found here that the hypermethylation resulted in further clustering of the affected genes and thus their expected movement towards a more repressive environment.

Since cancer cells are believed to experience more cell cycles than normal cells, we examined the chromatin structure at different stages in cell cycle for mouse, including G1, early S, late S to G2, and pre-M. Interestingly, we did observe an enhanced spatial separation between forest and prairie when the cell changes from G1 to early S stage, similar to what is observed in carcinogenesis (Figure 6A, 6B). Cells at early S stage possess lower F-F contact compared to F-P at genomic distances around 1M but the affected genes are significantly different from those affected in cancer cells by a similar structural chromatin change. The genes in the former process are significantly related to cell division, such as nucleosome assembly and DNA packaging (Supplementary Figure S8G). Cells from G1 to early S exhibit higher P-P contact than F-P at genomic distances around several million bases which is also similar to cancer cells. At large genomic distances, the chromosome structures of S-state cells are distinctly different from cancer cell lines. Forest domains in S-stage cells are seen to highly spatially segregate, consistent to a clustering of the early-replicating domain (*52, 53*). On the other hand, the long-range P domain aggregation is much weaker in S stage cells than in cancer cells. These observations further suggest that the chromatin structure change correlates with the change of biological functions and regulation in processes varying from carcinogenesis to mitosis, which presumably occur at very different in time scales. The similarity between structure changes (at the Mbp scale) of the two processes may suggest a role of cell division in cancer development.

**Figure 6.**
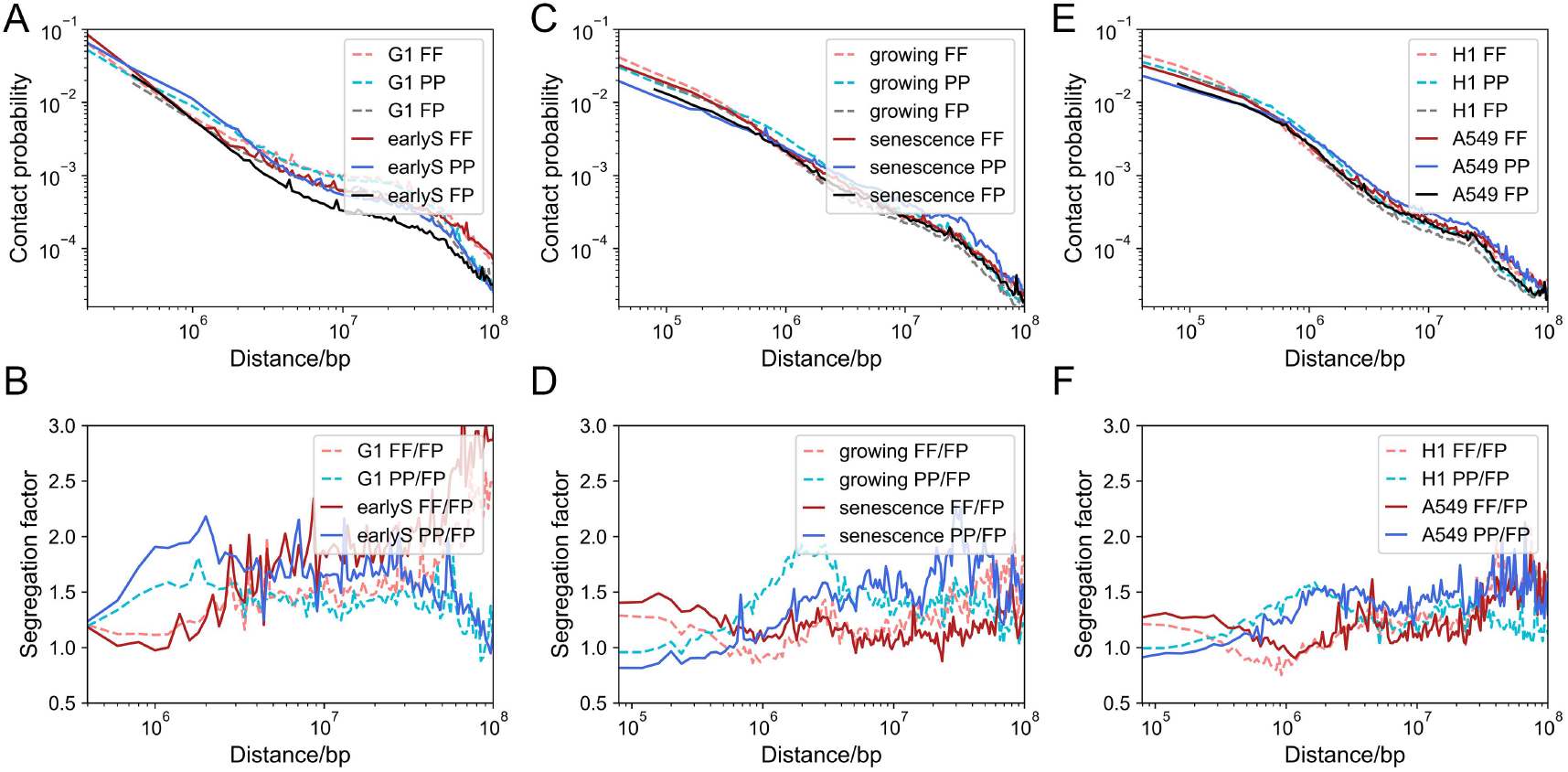
The Contact probability and segregation factor at varied genomic distances (chr1) for (A) (B) cells at G1 and early S stage in mouse cell cycle, (C) (D) growing cells and senescence cells, (E) (F) H1 and A549.

Cell senescence is also known to be highly influenced by cell cycles. It is a state of irreversible growth arrest, and is in certain sense a tumor suppressor. Senescence and carcinogenesis are mutually exclusive in most cases, although they can be induced by the same factors (*54, 55*). Interesting similarities do exist between cancer cell and senescence cell chromatin structures, such as enhanced long-range interactions, spatial segregation for repressive regions, analogous trend of hypomethylation of open sea (Figure 6C, 6D). The similar trend of increased domain segregation in both tumorigenesis and senescence suggest a common driving force shared by them and related to cell divisions. On the other hand, significant differences can also be observed between them. For CGI methylation, P-CGIs are more likely to be hypermethylated than F-CGIs in cancer cells while in aging cells, a larger portion of F-CGIs than that of P-CGIs are hypermethylated. In terms of chromatin structure, compared to growing cells, senescent cells lose and cancer cell lines gain local contacts for both forests and prairies. A higher portion of long-range chromatin contacts (especially that between forests and prairies) retained in the senescent than in the cancer cells. This latter difference may relate to cell identity retention, which is also a crucial difference between the highly and lowly-differentiated cancer cells.

Furthermore, important similarities can also be identified between early embryo development and carcinogenesis with respect to epigenetic regulation, gene expression, protein profiling and other important biological behaviors (*41*). From the chromatin structure point of view, short-range contact gains in the sacrifice of long-range ones are seen in both cancer cells and H1 (human embryonic stem cell line), in comparison to highly differentiated cells (Figure 6E, 6F). The former two are both characterized by high segregation factors at short genomic distances for forests and at long distances for prairies, although forests segregate more significantly at short distances and the prairie tends to cluster at longer distances in cancer cell lines than in H1.

The analyses performed here, in consistent with earlier studies, all show that different types of cancer appear to share similar overall changes of chromosomal structures, DNA methylation and gene expression, indicating the possibility of a rather general mechanism leading to carcinogenesis, especially from the structural perspective. Since almost no methylation changes are observed in CGI and open sea at very early stage of cancer, it is tempting to assume that methylation changes might not be the earliest changes in carcinogenesis. Except for epigenetic modifications, the expression of genes appears to also be influenced by sequence features at the CGI and forest/prairie scales. On the one hand, the local sequence of gene factors is crucial in that CGI genes and nonCGI genes display distinctly different expression activities. On the other hand, the context in which the gene resides is also important. The expression level of genes located in forests is generally higher than in those prairies and the former are more likely to be upregulated in carcinogenesis. A unified understanding of these epigenetic and transcriptional changes might find usefulness in diagnosis, or even treatment of cancers.

We note here that although our findings of chromatin structure changes during carcinogenesis presented in this paper are based on published results on cancer cell lines, our analyses on primary samples yielded similar results for both lung cancers and leukemia. These data will be presented in details elsewhere. In summary, the formations and changes of compartments, TAD, DNA methylation and gene expression all depend on the DNA sequence hierarchically: CGI or non-CGI locally, and sequence environment like CGI forest or prairie. Combining with the differences in gene density (especially housekeeping gene), gene tissue specificity, epigenetic marks, and transcription factor binding sites between forest and prairie discussed in our previous work, we propose that the genetic sequence itself provides the basic rule for the formation of high-order structure. As responses to the cell environment, structural modifiers, such as TFs, miRNA, DNA methyltransferase, histone modifiers, etc., all help to form specific chromosome structure and achieve cell identity and cell function. We leave the detailed discussion of these factors to a separate paper. We hope the heterogeneity of sequence could provide us with a new perspective to better understand various biological process intrinsically and help us develop more general cancer therapies.

**Supplemental Information** is available in the online version of the paper.

## Acknowledgement

We thank National Natural Science Foundation of China (21821004, 21873007) and National Key R&D Program of China (2017YFA0204702) in support of our research.

The results shown here are part based upon data generated by the TCGA Research Network: https://www.cancer.gov/tcga.

