## Supplemental Figures for "Domain segregated 3D chromatin structure and segmented DNA methylation in carcinogenesis"

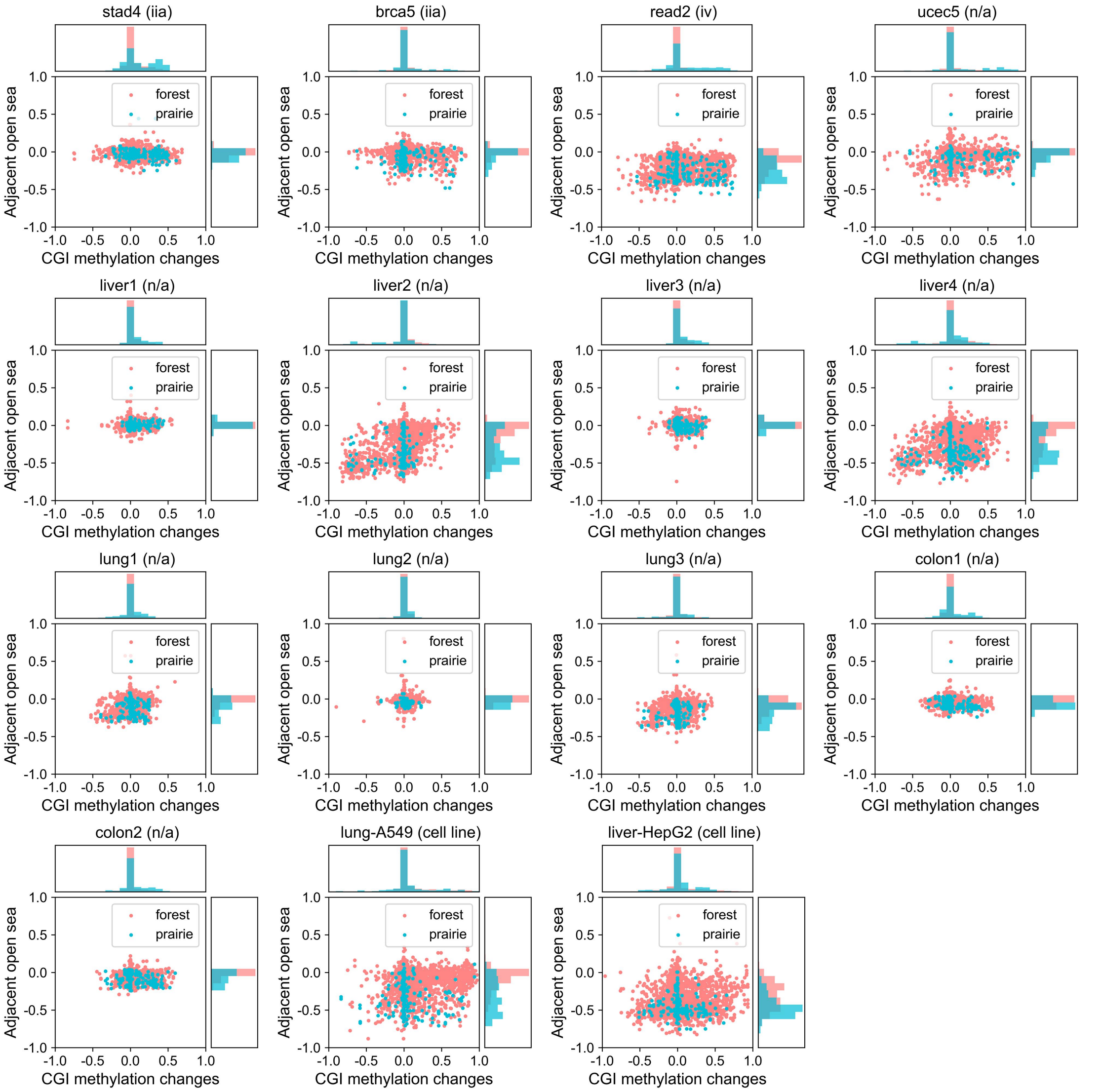


Supplementary Figure S1. Methylation changes in carcinogenesis. (A) Scatter plots for changes of methylation level in CGIs and open seas. Each dot represents the methylation changes of a CGI (x axis) and its adjacent open sea (y axis) from adjacent normal samples to cancer samples on chromosome 1. The probability density of changes of CGI and open sea are shown on the top and right sides of the figure, respectively.

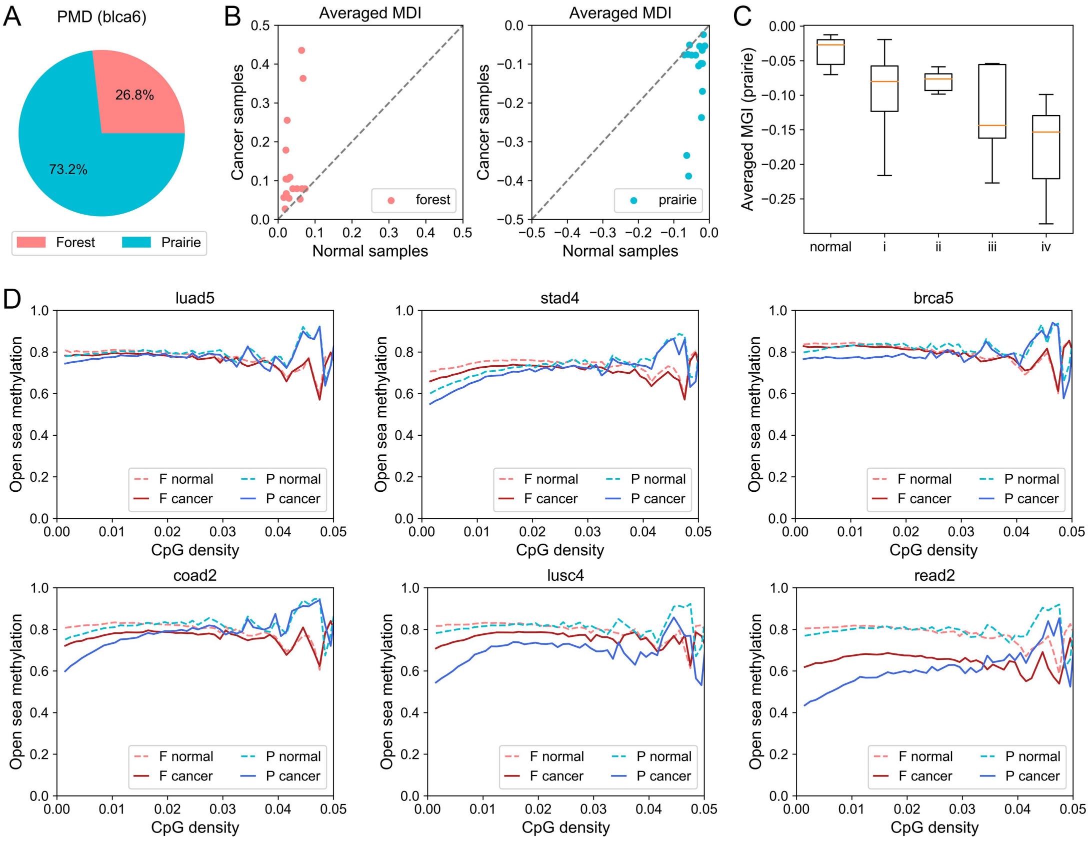


Supplementary Figure S2. (A) The sequence composition of PMD in blca6 cancer sample. Among all cancer samples, 72.9% of PMDs are located in prairie on average. (B) Averaged MDIs for forests and prairies in normal and cancer samples. Each dot represents one pair of samples. (C) The averaged MGIs of all prairie domains in normal samples and cancer samples in different stages. (D) CpG density and methylation level are calculated for all 1-kb beads in open sea. Each point on the curve shows the averaged methylation level for beads possess a given CpG density.

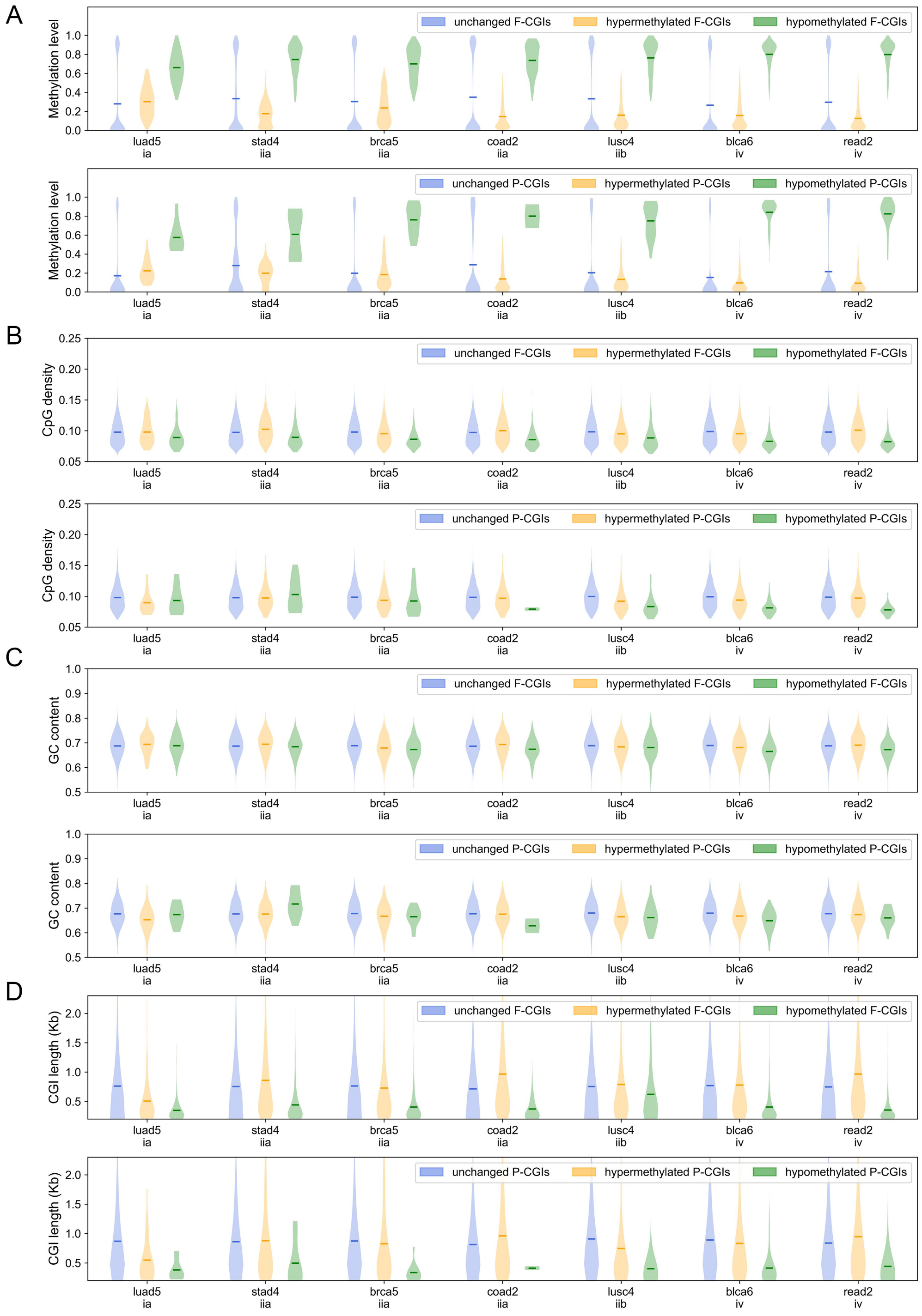


Supplementary Figure S3. (A) The methylation level, (B) CpG density, (C) GC content and (D) length for methylation unchanged CGI (uCGI), hypermethylated CGI (hyper-CGI) and hypomethylated CGI (hypo-CGI), respectively. Average level and *P*-value calculated by Welch's unequal variance t-test are shown in Supplementary Table S3.

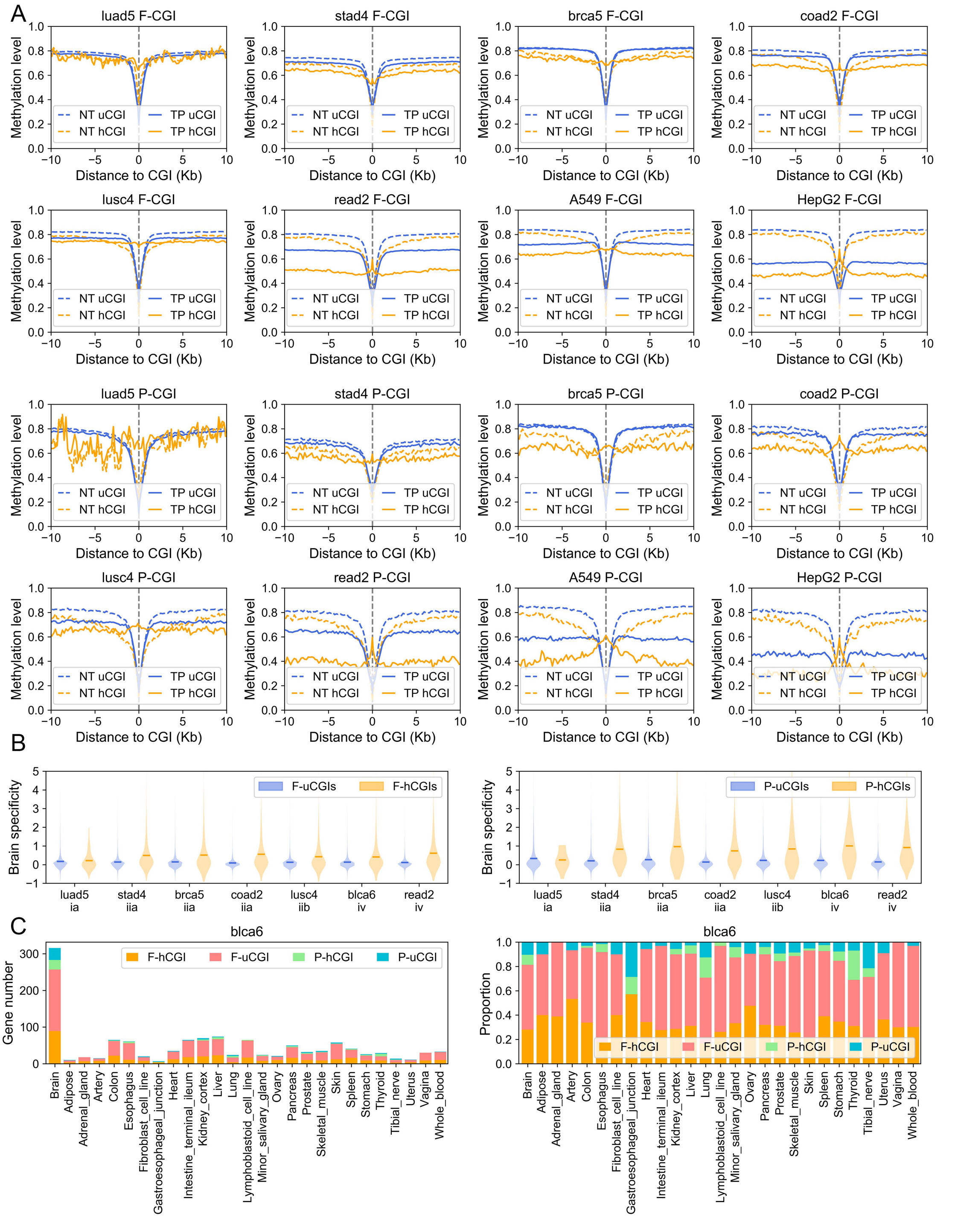


Supplementary Figure S4. (A) Methylation level of CGIs (at x=0) and their upstream (x<0) and downstream (x>0) CpGs. NT and TP refer to normal tissue and primary tumor, respectively. (B) The brain tissue specificity for uCGI and hCGI genes, *P*-values that calculated by Welch's unequal variance t-test are shown in Supplementary Table S4. (C) The number (left) and proportion (right) of F-uCGI, F-hCGI, P-uCGI and P-hCGI genes in tissue specific genes for different tissues.

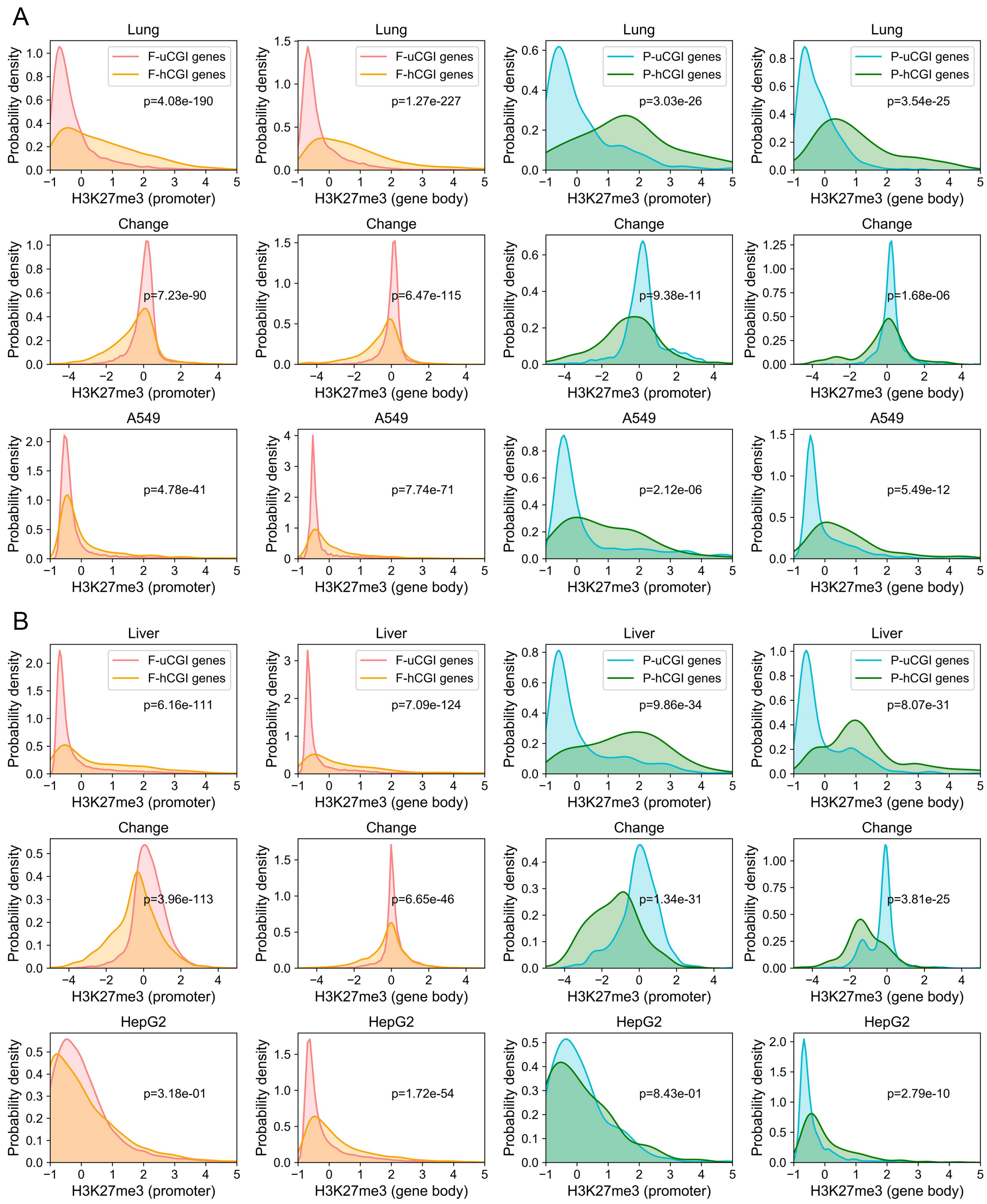


Supplementary Figure S5. The probability density of H3K27me3 for (A) lung tissue and cancer cell line A549 and (B) liver tissue and cancer cell line HepG2. The *P*-values that calculated by Welch's unequal variance t-test are shown on the figure.

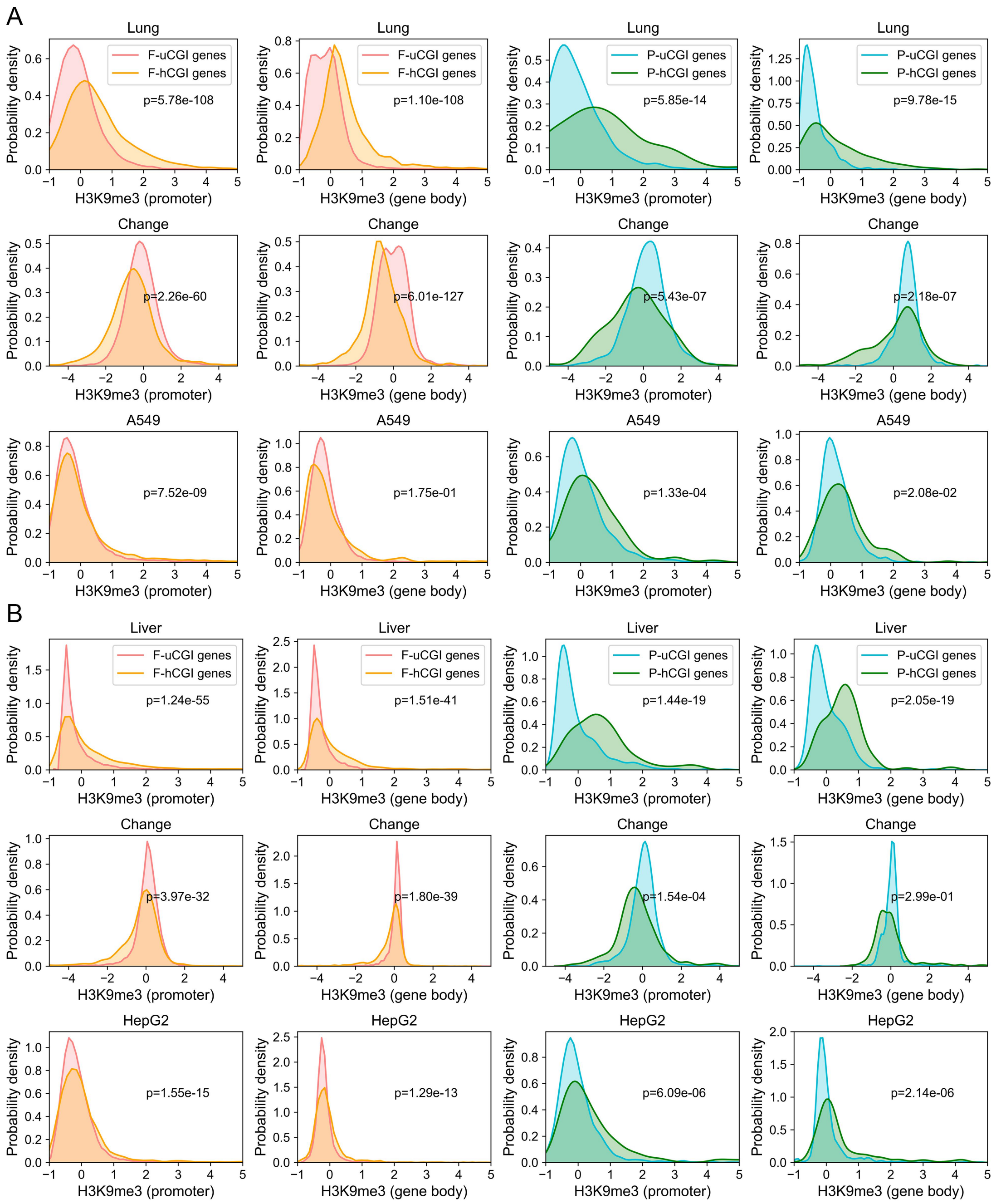


Supplementary Figure S6. The probability density of H3K9me3 for (A) lung tissue and cancer cell line A549 and (B) liver tissue and cancer cell line HepG2. The *P*-values that calculated by Welch's unequal variance t-test are shown on the figure.

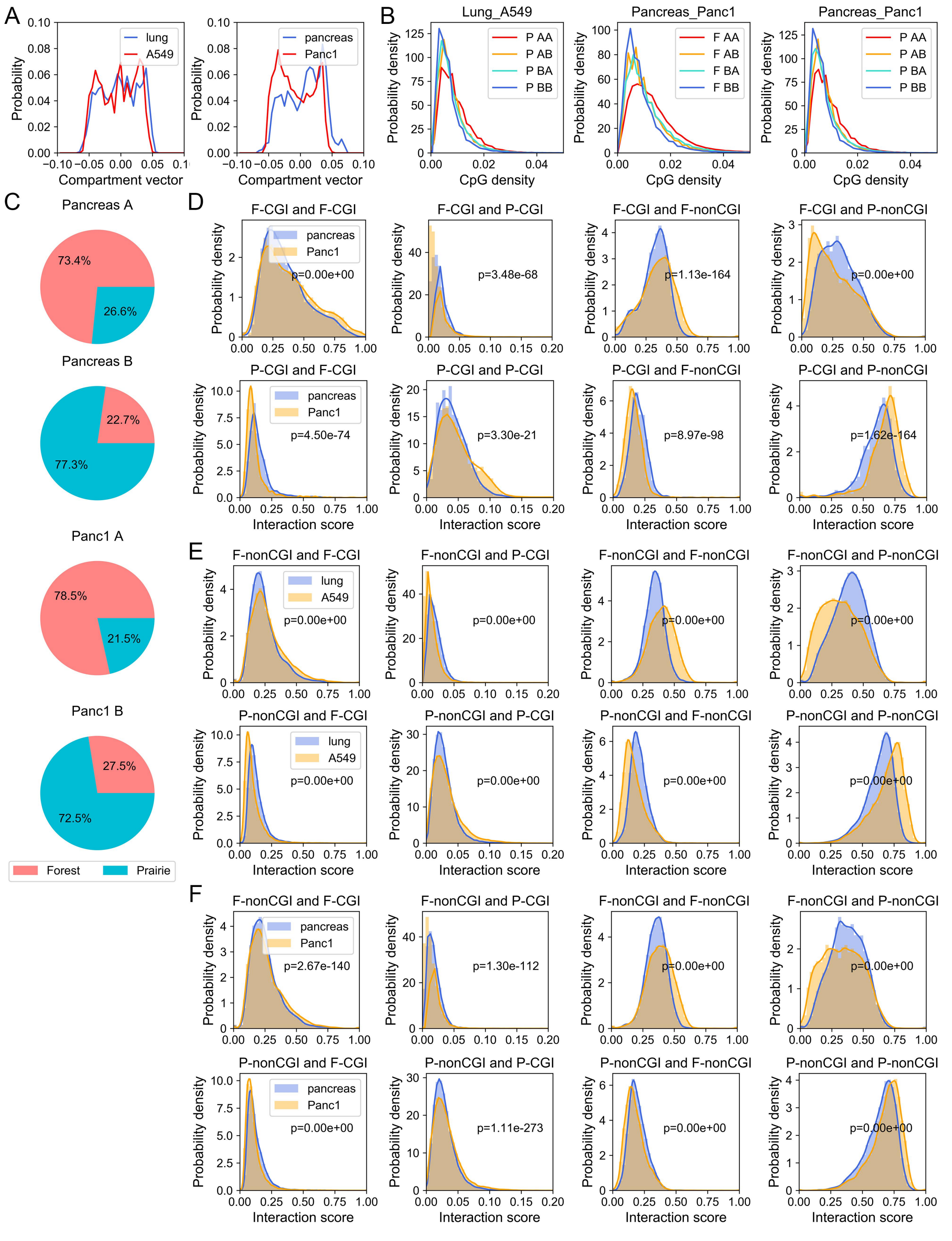

Supplementary Figure S7. The compartmentalization and CGI aggregation in carcinogenesis. (A) The probability distribution of compartment vector for lung and A549 (left), pancreas and Panc1 (right) on chromosome 1. (B) The probability density of CpG density for forest and prairie domains which belong to conservative compartment A (F AA) or B (F BB), as well as domains switch from A to B (F AB) or B to A (F BA) in carcinogenesis. (C) The proportion of forest and prairie sequences in compartments A and B for pancreas and Panc1. (D) The interaction scores between F-CGI (top) or P-CGI (bottom) and the four types of domians (F-CGI, F-nonCGI, P-CGI, and P-nonCGI) in normal pancreas and Panc1. Interaction scores for F-nonCGI and P-nonCGI are shown in (E) and (F). The *P*-values calculated by Welch's unequal variance t-test are shown on the figure and p=0 means that *P*-value < $10^{-300}$.

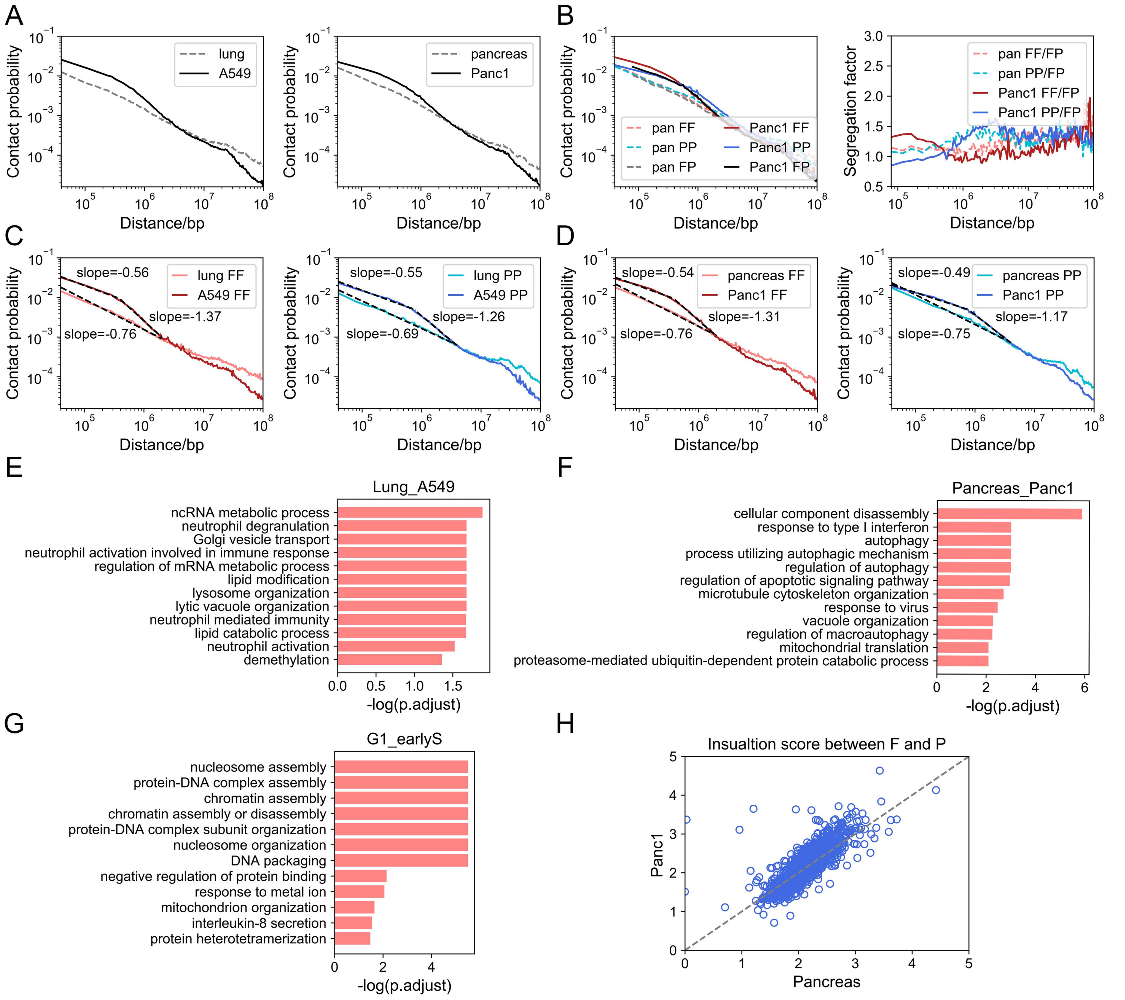


Supplementary Figure S8. General chromatin architecture in cancer cell lines. (A) The contact probability at varied genomic distances (chromosome 1 is used as an example). (B) The contact probability between forests and forests (FF), prairies and prairies (PP), forests and prairies (FP) at varied genomic distances for pancreas and Panc1. (C) The contact probability between forests (left) and between prairies (right) in lung and A549, as well as in (D) pancreas and Panc1. (E) GO analysis for forest genes which loss contact with other forest domains at 600K to 2M from lung to A549, (F) from pancreas to Panc1 and (G) from G1 to early S stage in mouse cell cycle. (H) The insulation score between adjacent forest and prairie domains in pancreas and Panc1.

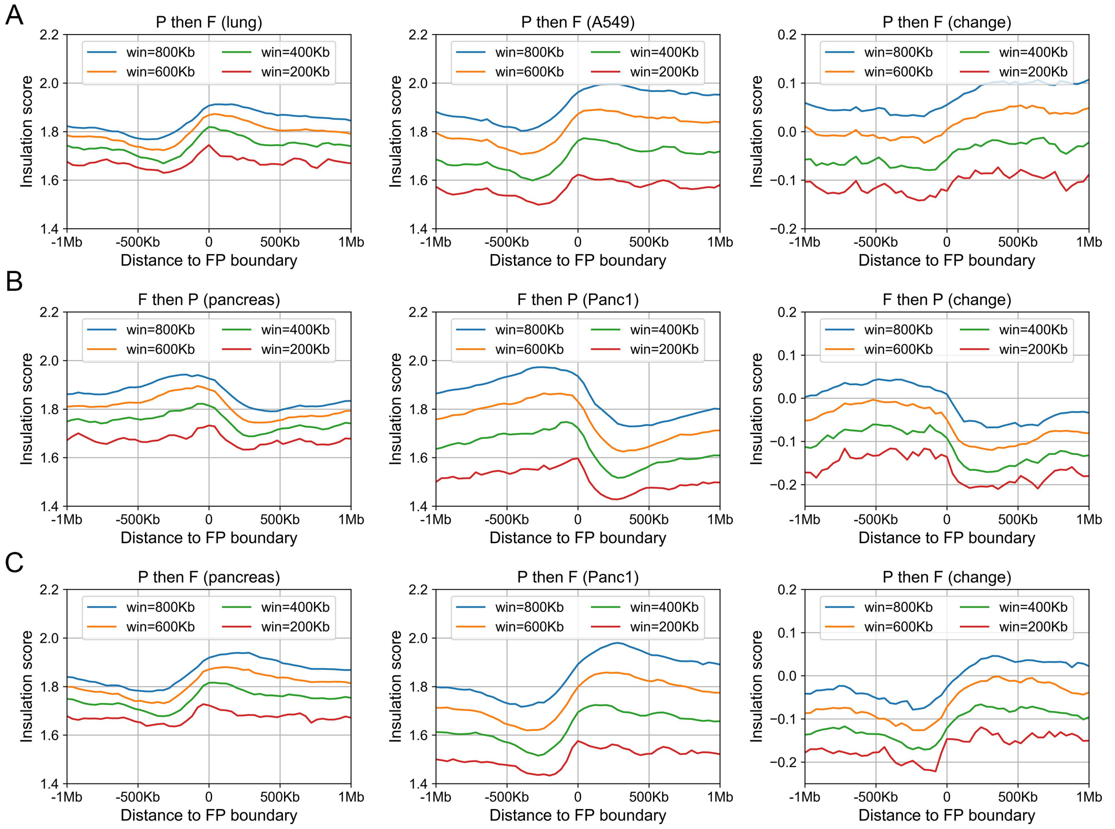


Supplementary Figure S9. (A) The insulation scores for 40-kb beads around P-F boundary at different window sizes in pancreas (left), Panc1 (middle) and the difference between A549 and lung (right). The data are aligned so that the prairie domain is left to the boundary (value 0). (B) and (C) show the insulation scores around F-P and P-F boundary, respectively, in pancreas and Panc1.

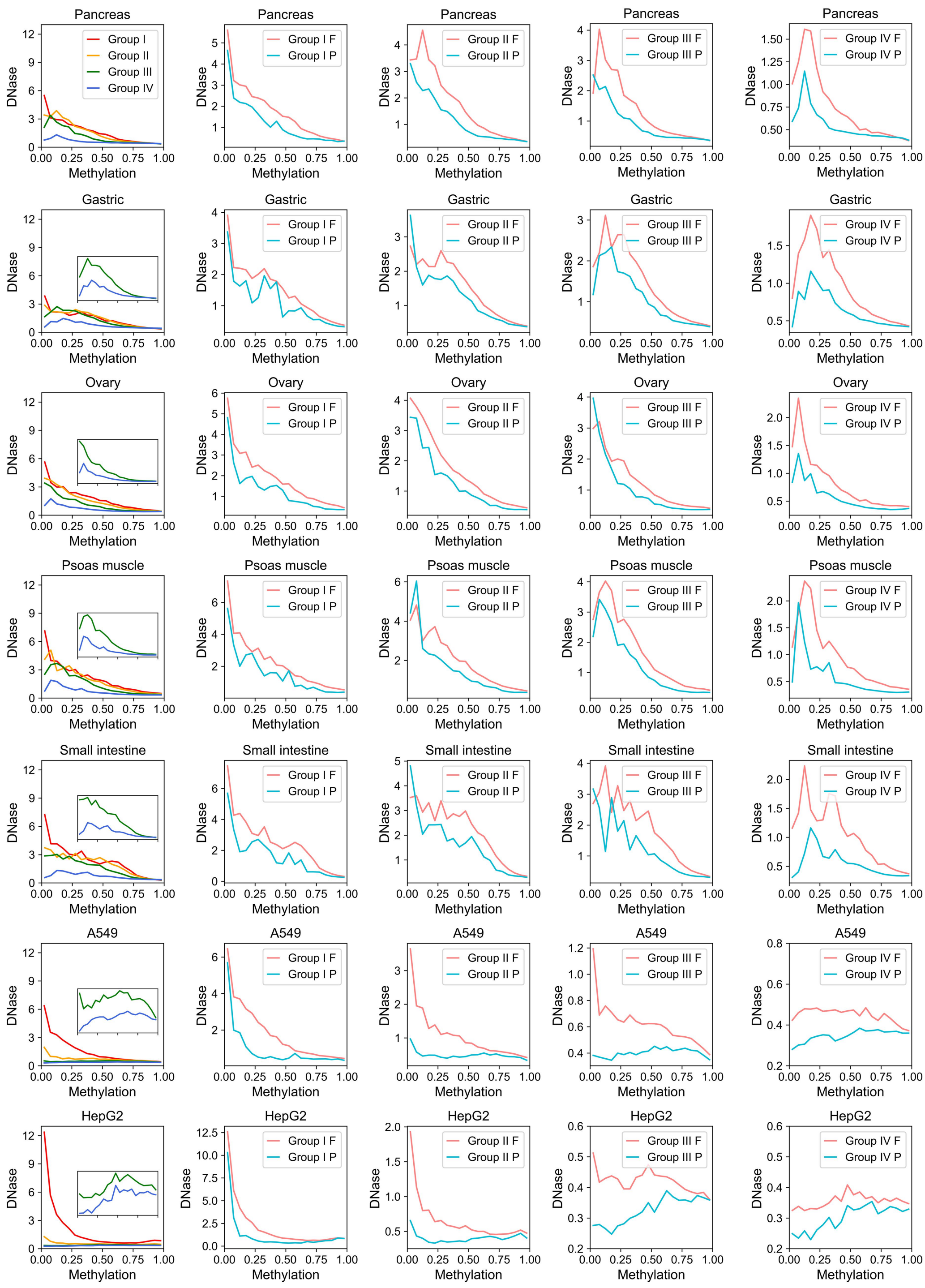


Supplementary Figure S10. Average DNase signal at various methylation levels. CpG density and CpG methylation level are calculated at a 1-kb resolution. Bins are divided to groups I, II, III or IV according to their CpG density (2.0%, 20.1%], (1.0%, 2.0%], (0.5%, 1.0%] or (0, 0.5%], respectively.

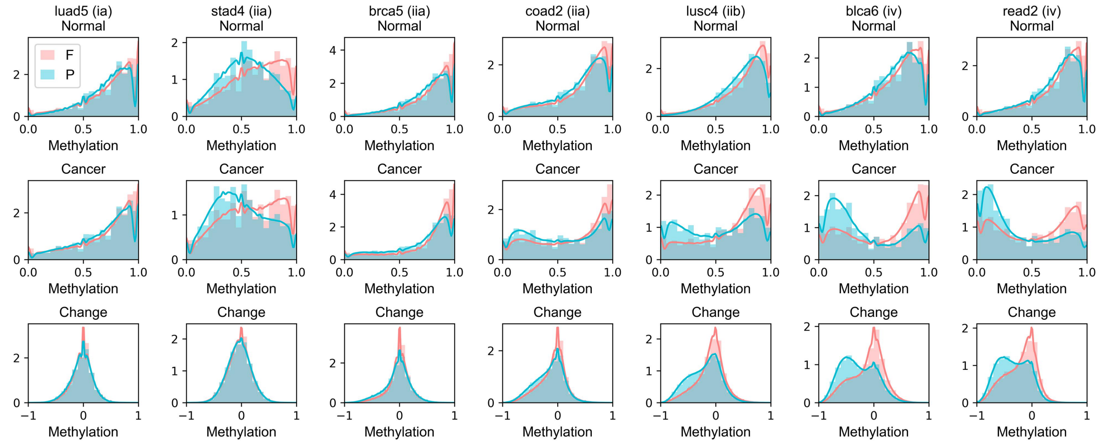


Supplementary Figure S11. The probability density of methylation level of solo-WCGW ('solo' refers to the CpGs with no neighboring CpGs and 'W' indicates A or T nucleotide). Each row is one sample. The methylation in normal tissue, cancer sample and changes are shown on the top, middle and bottom line, respectively.

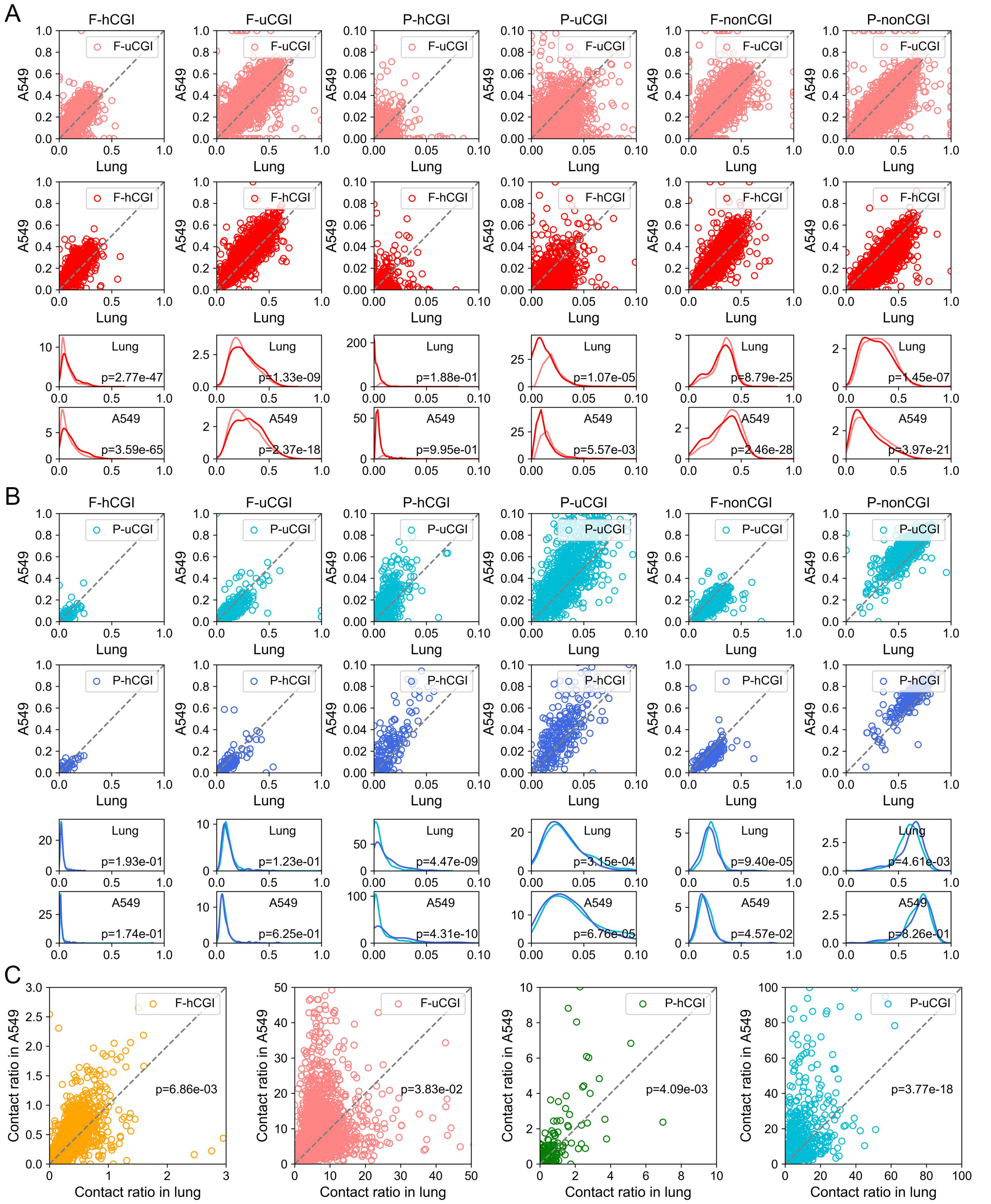


Supplementary Figure S12. The interaction scores for hCGIs and uCGIs. (A) The first row shows the interaction scores for F-hCGI with F-hCGI, F-uCGI, P-hCGI, P-uCGI, F-nonCGI and P-nonCGI (from left to right) in lung and A549. The second row shows the interaction scores for F-uCGI. The third and fourth row are the probability density of corresponding interaction scores in lung and A549, respectively. *P*-values are calculated by Welch's unequal variance t-test. (B) Interaction scores for P-hCGI and P-uCGI in lung and A549. (C) Contact ratio for F-hCGI, F-uCGI, P-hCGI and P-uCGI (from left to right). *P*-values are calculated by t-test.

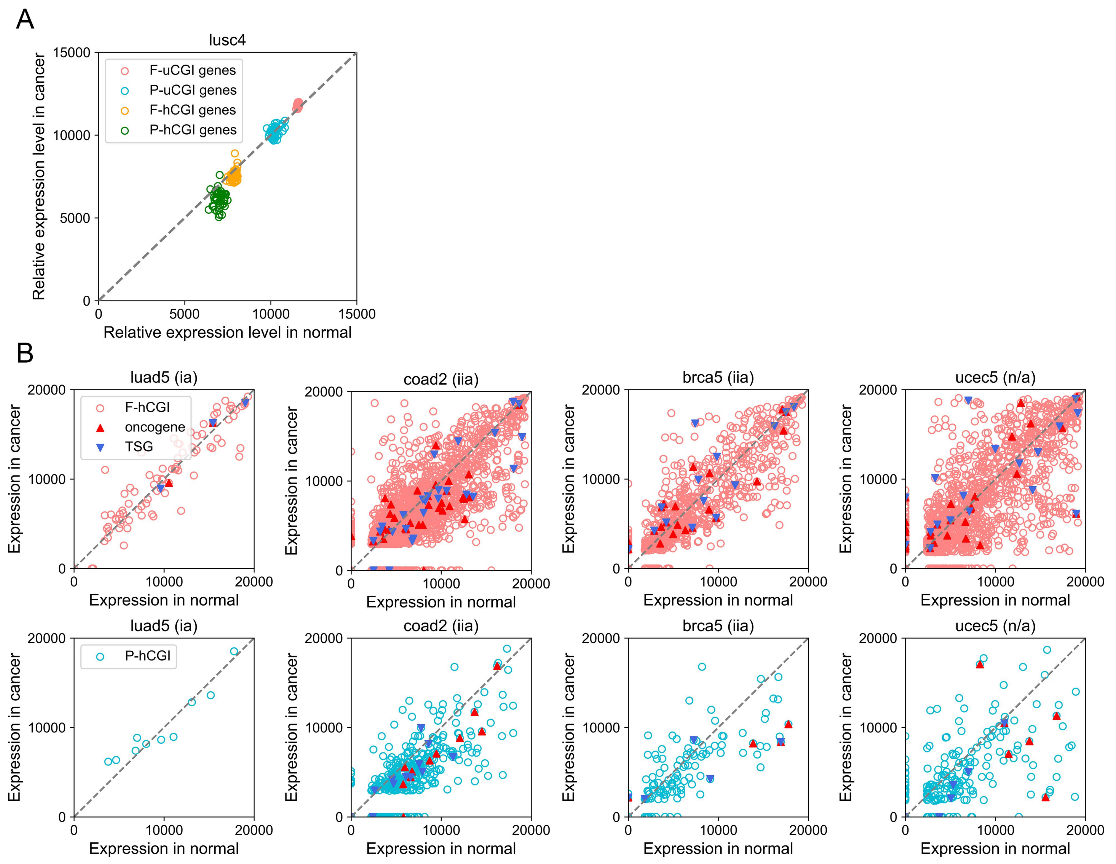


Supplementary Figure S13. (A) Difference of average relative expression level for hCGI and uCGI in 48 pairs of LUSC samples. (B) Expression level of F-hCGI genes (top) and P-hCGI genes (bottom) in four samples which have matching methylation and expression data. Each point represents one hCGI gene.
