## Supplemental Table 1 for "Domain segregated 3D chromatin structure and segmented DNA methylation in carcinogenesis"

### Supplemental Tables

Supplementary Table S1. Data information (RNA-seq data downloaded from TCGA)

| Sample ID | Cancer type | Sample ID | Cancer type |
|---|---|---|---|
| TCGA-BL-A13J | Bladder Urothelial Carcinoma (BLCA) | TCGA-BC-A10Q | Liver hepatocellular carcinoma (LIHC) |
| TCGA-BT-A20N | | TCGA-BC-A10R | |
| TCGA-BT-A20Q | | TCGA-BC-A10T | |
| TCGA-BT-A20R | | TCGA-BC-A10U | |
| TCGA-BT-A20U | | TCGA-BC-A10W | |
| TCGA-BT-A20W | | TCGA-BC-A10X | |
| TCGA-BT-A2LA | | TCGA-BC-A10Y | |
| TCGA-BT-A2LB | | TCGA-BC-A10Z | |
| TCGA-CU-A0YN | | TCGA-BC-A110 | |
| TCGA-CU-A0YR | | TCGA-BC-A216 | |
| TCGA-GC-A3BM | | TCGA-BD-A2L6 | |
| TCGA-GC-A3WC | | TCGA-BD-A3EP | |
| TCGA-GC-A6I3 | | TCGA-DD-A113 | |
| TCGA-GD-A2C5 | | TCGA-DD-A114 | |
| TCGA-GD-A3OP | | TCGA-DD-A116 | |
| TCGA-GD-A3OQ | | TCGA-DD-A118 | |
| TCGA-K4-A3WV | | TCGA-DD-A119 | |
| TCGA-K4-A54R | | TCGA-DD-A11A | |
| TCGA-K4-A5RI | | TCGA-DD-A11B | |
| TCGA-A7-A0CE | | TCGA-DD-A11C | |
| TCGA-A7-A0CH | | TCGA-DD-A11D | |
| TCGA-A7-A0D9 | | TCGA-DD-A1EB | |
| TCGA-A7-A0DB | | TCGA-DD-A1EC | |
| TCGA-A7-A0DC | | TCGA-DD-A1EE | |
| TCGA-A7-A13E | | TCGA-DD-A1EG | |
| TCGA-A7-A13F | | TCGA-DD-A1EH | |
| TCGA-AC-A23H | | TCGA-DD-A1EI | |
| TCGA-AC-A2FB | | TCGA-DD-A1EJ | |
| TCGA-AC-A2FF | | TCGA-DD-A1EL | |
| TCGA-BH-A0AU | | TCGA-DD-A39V | |
| TCGA-BH-A0AY | | TCGA-DD-A39W | |
| TCGA-BH-A0AZ | | TCGA-DD-A39X | |
| TCGA-BH-A0B5 | | TCGA-DD-A39Z | |
| TCGA-BH-A0B7 | | TCGA-DD-A3A1 | |
| TCGA-BH-A0B8 | | TCGA-DD-A3A2 | |
| TCGA-BH-A0BA | | TCGA-DD-A3A3 | |
| TCGA-BH-A0BC | | TCGA-DD-A3A4 | |
| TCGA-BH-A0BJ | | TCGA-DD-A3A5 | |
| TCGA-BH-A0BM | | TCGA-DD-A3A6 | |

| | | | |
|---|---|---|---|
| TCGA-BH-A0BQ | | TCGA-DD-A3A8 | |
| TCGA-BH-A0BT | | TCGA-EP-A12J | |
| TCGA-BH-A0BV | | TCGA-EP-A26S | |
| TCGA-BH-A0BW | | TCGA-EP-A3RK | |
| TCGA-BH-A0BZ | | TCGA-ES-A2HT | |
| TCGA-BH-A0C0 | | TCGA-FV-A23B | |
| TCGA-BH-A0C3 | | TCGA-FV-A2QR | |
| TCGA-BH-A0DD | | TCGA-FV-A3I0 | |
| TCGA-BH-A0DG | | TCGA-FV-A3I1 | |
| TCGA-BH-A0DH | | TCGA-FV-A3R2 | |
| TCGA-BH-A0DK | | TCGA-G3-A3CH | |
| TCGA-BH-A0DL | | TCGA-38-4625 | Lung adenocarcinoma (LUAD) |
| TCGA-BH-A0DP | | TCGA-38-4626 | |
| TCGA-BH-A0DQ | | TCGA-38-4627 | |
| TCGA-BH-A0DT | | TCGA-38-4632 | |
| TCGA-BH-A0DV | | TCGA-44-2655 | |
| TCGA-BH-A0DZ | | TCGA-44-2657 | |
| TCGA-BH-A0E0 | | TCGA-44-2661 | |
| TCGA-BH-A0E1 | | TCGA-44-2662 | |
| TCGA-BH-A0H5 | | TCGA-44-2665 | |
| TCGA-BH-A0H7 | | TCGA-44-2668 | |
| TCGA-BH-A0H9 | | TCGA-44-3396 | |
| TCGA-BH-A0HA | | TCGA-44-5645 | |
| TCGA-BH-A0HK | | TCGA-44-6145 | |
| TCGA-BH-A18J | | TCGA-44-6146 | |
| TCGA-BH-A18K | | TCGA-44-6147 | |
| TCGA-BH-A18L | | TCGA-44-6148 | |
| TCGA-BH-A18M | | TCGA-44-6776 | |
| TCGA-BH-A18N | | TCGA-44-6777 | |
| TCGA-BH-A18P | | TCGA-44-6778 | |
| TCGA-BH-A18Q | | TCGA-49-4490 | |
| TCGA-BH-A18R | | TCGA-49-4512 | |
| TCGA-BH-A18S | | TCGA-49-6742 | |
| TCGA-BH-A18U | | TCGA-49-6743 | |
| TCGA-BH-A18V | | TCGA-49-6744 | |
| TCGA-BH-A1EN | | TCGA-49-6745 | |
| TCGA-BH-A1EO | | TCGA-49-6761 | |
| TCGA-BH-A1EU | | TCGA-50-5930 | |
| TCGA-BH-A1EV | | TCGA-50-5931 | |
| TCGA-BH-A1F2 | | TCGA-50-5932 | |
| TCGA-BH-A1FB | | TCGA-50-5933 | |
| TCGA-BH-A1FC | | TCGA-50-5935 | |
| TCGA-BH-A1FN | | TCGA-50-5936 | |

| | | | |
|---|---|---|---|
| TCGA-BH-A1FU | | TCGA-50-5939 | |
| TCGA-BH-A203 | | TCGA-50-6595 | |
| TCGA-BH-A204 | | TCGA-55-6968 | |
| TCGA-BH-A208 | | TCGA-55-6970 | |
| TCGA-BH-A209 | | TCGA-55-6971 | |
| TCGA-E2-A153 | | TCGA-55-6972 | |
| TCGA-E2-A158 | | TCGA-55-6975 | |
| TCGA-E2-A15I | | TCGA-55-6978 | |
| TCGA-E2-A15K | | TCGA-55-6979 | |
| TCGA-E2-A15M | | TCGA-55-6980 | |
| TCGA-E2-A1BC | | TCGA-55-6981 | |
| TCGA-E2-A1IG | | TCGA-55-6982 | |
| TCGA-E2-A1L7 | | TCGA-55-6983 | |
| TCGA-E2-A1LB | | TCGA-55-6984 | |
| TCGA-E2-A1LH | | TCGA-55-6985 | |
| TCGA-E2-A1LS | | TCGA-55-6986 | |
| TCGA-E9-A1N4 | | TCGA-73-4676 | |
| TCGA-E9-A1N5 | | TCGA-91-6828 | |
| TCGA-E9-A1N6 | | TCGA-91-6829 | |
| TCGA-E9-A1N9 | | TCGA-91-6831 | |
| TCGA-E9-A1NA | | TCGA-91-6835 | |
| TCGA-E9-A1ND | | TCGA-91-6836 | |
| TCGA-E9-A1NF | | TCGA-91-6847 | |
| TCGA-E9-A1NG | | TCGA-91-6849 | |
| TCGA-E9-A1R7 | | TCGA-22-4593 | Lung squamous cell carcinoma (LUSC) |
| TCGA-E9-A1RB | | TCGA-22-4609 | |
| TCGA-E9-A1RC | | TCGA-22-5471 | |
| TCGA-E9-A1RD | | TCGA-22-5472 | |
| TCGA-E9-A1RF | | TCGA-22-5478 | |
| TCGA-E9-A1RH | | TCGA-22-5481 | |
| TCGA-E9-A1RI | | TCGA-22-5482 | |
| TCGA-GI-A2C8 | | TCGA-22-5483 | |
| TCGA-GI-A2C9 | | TCGA-22-5489 | |
| TCGA-FU-A3EO | Cervical squamous cell carcinoma and endocervical adenocarcinoma (CESC) | TCGA-22-5491 | |
| TCGA-HM-A3JJ | | TCGA-33-4587 | |
| TCGA-MY-A5BF | | TCGA-33-6737 | |
| TCGA-W5-AA2I | Cholangiocarcinoma (CHOL) | TCGA-34-7107 | |
| TCGA-W5-AA2Q | | TCGA-34-8454 | |
| TCGA-W5-AA2R | | TCGA-39-5040 | |
| TCGA-W5-AA2U | | TCGA-43-5670 | |
| TCGA-W5-AA2X | | TCGA-43-6143 | |
| TCGA-W5-AA30 | | TCGA-43-6647 | |

| | | | |
|---|---|---|---|
| TCGA-W5-AA31 | | TCGA-43-6771 | |
| TCGA-W5-AA34 | | TCGA-43-6773 | |
| TCGA-ZU-A8S4 | | TCGA-43-7657 | |
| TCGA-A6-2671 | Colon adenocarcinoma (COAD) | TCGA-43-7658 | |
| TCGA-A6-2675 | | TCGA-51-4079 | |
| TCGA-A6-2678 | | TCGA-51-4081 | |
| TCGA-A6-2679 | | TCGA-56-7222 | |
| TCGA-A6-2680 | | TCGA-56-7579 | |
| TCGA-A6-2682 | | TCGA-56-7580 | |
| TCGA-A6-2683 | | TCGA-56-7582 | |
| TCGA-A6-2684 | | TCGA-56-7730 | |
| TCGA-A6-2685 | | TCGA-56-7731 | |
| TCGA-A6-2686 | | TCGA-56-8082 | |
| TCGA-A6-5659 | | TCGA-56-8083 | |
| TCGA-A6-5662 | | TCGA-56-8201 | |
| TCGA-A6-5665 | | TCGA-56-8309 | |
| TCGA-A6-5667 | | TCGA-56-8623 | |
| TCGA-AA-3489 | | TCGA-58-8386 | |
| TCGA-AA-3496 | | TCGA-60-2709 | |
| TCGA-AA-3511 | | TCGA-77-7138 | |
| TCGA-AA-3514 | | TCGA-77-7142 | |
| TCGA-AA-3516 | | TCGA-77-7335 | |
| TCGA-AA-3517 | | TCGA-77-7337 | |
| TCGA-AA-3518 | | TCGA-77-7338 | |
| TCGA-AA-3520 | | TCGA-77-8007 | |
| TCGA-AA-3522 | | TCGA-77-8008 | |
| TCGA-AA-3525 | | TCGA-85-7710 | |
| TCGA-AA-3527 | | TCGA-90-6837 | |
| TCGA-AA-3531 | | TCGA-90-7767 | |
| TCGA-AA-3534 | | TCGA-92-7340 | |
| TCGA-AA-3655 | | TCGA-H6-8124 | Pancreatic adenocarcinoma (PAAD) |
| TCGA-AA-3660 | | TCGA-H6-A45N | |
| TCGA-AA-3662 | | TCGA-HV-A5A3 | |
| TCGA-AA-3663 | | TCGA-YB-A89D | |
| TCGA-AA-3697 | | TCGA-P8-A5KC | Pheochromocytoma and Paraganglioma (PCPG) |
| TCGA-AA-3712 | | TCGA-P8-A5KD | |
| TCGA-AA-3713 | | TCGA-SQ-A6I4 | |
| TCGA-AZ-6598 | | TCGA-CH-5761 | Prostate adenocarcinoma (PRAD) |
| TCGA-AZ-6599 | | TCGA-CH-5768 | |
| TCGA-AZ-6600 | | TCGA-CH-5769 | |
| TCGA-AZ-6601 | | TCGA-EJ-7115 | |
| TCGA-AZ-6603 | | TCGA-EJ-7123 | |
| TCGA-AZ-6605 | | TCGA-EJ-7125 | |

| | | |
|---|---|---|
| TCGA-F4-6704 | | TCGA-EJ-7314 |
| TCGA-IC-A6RE | Esophageal carcinoma (ESCA) | TCGA-EJ-7315 |
| TCGA-IC-A6RF | | TCGA-EJ-7317 |
| TCGA-L5-A43C | | TCGA-EJ-7321 |
| TCGA-L5-A4OG | | TCGA-EJ-7327 |
| TCGA-L5-A4OJ | | TCGA-EJ-7328 |
| TCGA-L5-A4OO | | TCGA-EJ-7330 |
| TCGA-V5-A7RE | | TCGA-EJ-7331 |
| TCGA-V5-AASX | | TCGA-EJ-7781 |
| TCGA-CV-6933 | Head and Neck squamous cell carcinoma (HNSC) | TCGA-EJ-7782 |
| TCGA-CV-6934 | | TCGA-EJ-7783 |
| TCGA-CV-6935 | | TCGA-EJ-7784 |
| TCGA-CV-6936 | | TCGA-EJ-7785 |
| TCGA-CV-6938 | | TCGA-EJ-7786 |
| TCGA-CV-6939 | | TCGA-EJ-7789 |
| TCGA-CV-6943 | | TCGA-EJ-7792 |
| TCGA-CV-6955 | | TCGA-EJ-7793 |
| TCGA-CV-6956 | | TCGA-EJ-7794 |
| TCGA-CV-6959 | | TCGA-EJ-7797 |
| TCGA-CV-6960 | | TCGA-EJ-A8FO |
| TCGA-CV-6961 | | TCGA-G9-6333 |
| TCGA-CV-6962 | | TCGA-G9-6342 |
| TCGA-CV-7091 | | TCGA-G9-6348 |
| TCGA-CV-7097 | | TCGA-G9-6351 |
| TCGA-CV-7101 | | TCGA-G9-6356 |
| TCGA-CV-7103 | | TCGA-G9-6362 |
| TCGA-CV-7177 | | TCGA-G9-6363 |
| TCGA-CV-7178 | | TCGA-G9-6365 |
| TCGA-CV-7183 | | TCGA-G9-6384 |
| TCGA-CV-7235 | | TCGA-G9-6496 |
| TCGA-CV-7238 | | TCGA-G9-6499 |
| TCGA-CV-7242 | | TCGA-HC-7211 |
| TCGA-CV-7245 | | TCGA-HC-7737 |
| TCGA-CV-7250 | | TCGA-HC-7738 |
| TCGA-CV-7252 | | TCGA-HC-7740 |
| TCGA-CV-7255 | | TCGA-HC-7742 |
| TCGA-CV-7261 | | TCGA-HC-7745 |
| TCGA-CV-7406 | | TCGA-HC-7747 |
| TCGA-CV-7416 | | TCGA-HC-7752 |
| TCGA-CV-7423 | | TCGA-HC-7819 |
| TCGA-CV-7424 | | TCGA-HC-8258 |
| TCGA-CV-7425 | | TCGA-HC-8259 |
| TCGA-CV-7432 | | TCGA-HC-8260 |

| | | | |
|---|---|---|---|
| TCGA-CV-7434 | | TCGA-HC-8262 | |
| TCGA-CV-7437 | | TCGA-J4-A83J | |
| TCGA-CV-7438 | | TCGA-AF-2691 | Rectum adenocarcinoma (READ) |
| TCGA-CV-7440 | | TCGA-AF-2692 | |
| TCGA-H7-A6C4 | | TCGA-AF-3400 | |
| TCGA-HD-8635 | | TCGA-AF-5654 | |
| TCGA-HD-A6HZ | | TCGA-AG-3725 | |
| TCGA-HD-A6I0 | | TCGA-AG-3731 | |
| TCGA-WA-A7GZ | | TCGA-AG-3732 | |
| TCGA-KL-8324 | Kidney Chromophobe (KICH) | TCGA-AG-3742 | |
| TCGA-KL-8326 | | TCGA-AH-6643 | |
| TCGA-KL-8329 | | TCGA-FX-A2QS | Sarcoma (SARC) |
| TCGA-KL-8336 | | TCGA-K1-A3PO | |
| TCGA-KL-8339 | | TCGA-BR-6453 | Stomach adenocarcinoma (STAD) |
| TCGA-KN-8419 | | TCGA-BR-6454 | |
| TCGA-KN-8423 | | TCGA-BR-6457 | |
| TCGA-KN-8424 | | TCGA-BR-6802 | |
| TCGA-KN-8425 | | TCGA-BR-7704 | |
| TCGA-KN-8426 | | TCGA-BR-7715 | |
| TCGA-KN-8427 | | TCGA-BR-7716 | |
| TCGA-KN-8428 | | TCGA-BR-7717 | |
| TCGA-KN-8429 | | TCGA-BR-7851 | |
| TCGA-KN-8430 | | TCGA-BR-8060 | |
| TCGA-KN-8431 | | TCGA-CG-5720 | |
| TCGA-KN-8432 | | TCGA-CG-5721 | |
| TCGA-KN-8433 | | TCGA-CG-5722 | |
| TCGA-KN-8434 | | TCGA-CG-5734 | |
| TCGA-KN-8435 | | TCGA-FP-7735 | |
| TCGA-KN-8436 | | TCGA-FP-7829 | |
| TCGA-KN-8437 | | TCGA-HU-8238 | |
| TCGA-KO-8403 | | TCGA-HU-A4GC | |
| TCGA-KO-8415 | | TCGA-HU-A4GH | |
| TCGA-A3-3358 | Kidney renal clear cell carcinoma (KIRC) | TCGA-HU-A4GP | |
| TCGA-A3-3387 | | TCGA-HU-A4GY | |
| TCGA-B0-4700 | | TCGA-HU-A4HB | |
| TCGA-B0-4712 | | TCGA-IN-7806 | |
| TCGA-B0-5402 | | TCGA-IN-8663 | |
| TCGA-B0-5690 | | TCGA-IN-AB1V | |
| TCGA-B0-5691 | | TCGA-IN-AB1X | |
| TCGA-B0-5694 | | TCGA-IP-7968 | |
| TCGA-B0-5696 | | TCGA-BJ-A28R | Thyroid carcinoma (THCA) |
| TCGA-B0-5697 | | TCGA-BJ-A28W | |
| TCGA-B0-5699 | | TCGA-BJ-A28X | |

| | | |
|---|---|---|
| TCGA-B0-5701 | | TCGA-BJ-A290 |
| TCGA-B0-5703 | | TCGA-BJ-A2N7 |
| TCGA-B0-5705 | | TCGA-BJ-A2N8 |
| TCGA-B0-5706 | | TCGA-BJ-A2N9 |
| TCGA-B0-5709 | | TCGA-BJ-A2NA |
| TCGA-B0-5711 | | TCGA-BJ-A3PR |
| TCGA-B0-5712 | | TCGA-BJ-A3PU |
| TCGA-B2-5636 | | TCGA-DO-A1JZ |
| TCGA-B2-5641 | | TCGA-E8-A2JQ |
| TCGA-B8-4619 | | TCGA-EL-A3GZ |
| TCGA-B8-4620 | | TCGA-EL-A3H1 |
| TCGA-B8-4622 | | TCGA-EL-A3H2 |
| TCGA-B8-5549 | | TCGA-EL-A3H7 |
| TCGA-CJ-5672 | | TCGA-EL-A3MW |
| TCGA-CJ-5676 | | TCGA-EL-A3MX |
| TCGA-CJ-5677 | | TCGA-EL-A3MY |
| TCGA-CJ-5678 | | TCGA-EL-A3N2 |
| TCGA-CJ-5679 | | TCGA-EL-A3N3 |
| TCGA-CJ-5680 | | TCGA-EL-A3T0 |
| TCGA-CJ-5681 | | TCGA-EL-A3T1 |
| TCGA-CJ-5689 | | TCGA-EL-A3T2 |
| TCGA-CJ-6030 | | TCGA-EL-A3T3 |
| TCGA-CJ-6033 | | TCGA-EL-A3T6 |
| TCGA-CW-5580 | | TCGA-EL-A3T7 |
| TCGA-CW-5581 | | TCGA-EL-A3T8 |
| TCGA-CW-5584 | | TCGA-EL-A3TA |
| TCGA-CW-5585 | | TCGA-EL-A3TB |
| TCGA-CW-5587 | | TCGA-EL-A3ZG |
| TCGA-CW-5589 | | TCGA-EL-A3ZH |
| TCGA-CW-5591 | | TCGA-EL-A3ZK |
| TCGA-CW-6087 | | TCGA-EL-A3ZL |
| TCGA-CW-6088 | | TCGA-EL-A3ZM |
| TCGA-CW-6090 | | TCGA-EL-A3ZO |
| TCGA-CZ-4863 | | TCGA-EL-A3ZP |
| TCGA-CZ-4864 | | TCGA-EL-A3ZQ |
| TCGA-CZ-4865 | | TCGA-EL-A3ZR |
| TCGA-CZ-5451 | | TCGA-EL-A3ZS |
| TCGA-CZ-5452 | | TCGA-EL-A3ZT |
| TCGA-CZ-5453 | | TCGA-EM-A1CS |
| TCGA-CZ-5454 | | TCGA-EM-A1CT |
| TCGA-CZ-5455 | | TCGA-EM-A1CU |
| TCGA-CZ-5456 | | TCGA-EM-A1CV |
| TCGA-CZ-5457 | | TCGA-EM-A1CW |

| | | | |
|---|---|---|---|
| TCGA-CZ-5458 | | TCGA-EM-A1YC | |
| TCGA-CZ-5461 | | TCGA-EM-A3ST | |
| TCGA-CZ-5462 | | TCGA-ET-A3DP | |
| TCGA-CZ-5463 | | TCGA-ET-A3DW | |
| TCGA-CZ-5465 | | TCGA-FY-A3TY | |
| TCGA-CZ-5466 | | TCGA-GE-A2C6 | |
| TCGA-CZ-5467 | | TCGA-H2-A2K9 | |
| TCGA-CZ-5468 | | TCGA-KS-A41I | |
| TCGA-CZ-5469 | | TCGA-KS-A41J | |
| TCGA-CZ-5470 | | TCGA-KS-A41L | |
| TCGA-CZ-5982 | | TCGA-X7-A8D6 | Thymoma (THYM) |
| TCGA-CZ-5984 | | TCGA-X7-A8D7 | |
| TCGA-CZ-5985 | | TCGA-AJ-A2QL | Uterine Corpus Endometrial Carcinoma (UCEC) |
| TCGA-CZ-5986 | | TCGA-AJ-A3NC | |
| TCGA-CZ-5987 | | TCGA-AJ-A3NE | |
| TCGA-CZ-5988 | | TCGA-AJ-A3NH | |
| TCGA-CZ-5989 | | TCGA-AX-A05Y | |
| TCGA-A4-A4ZT | Kidney renal papillary cell carcinoma (KIRP) | TCGA-AX-A0IZ | |
| TCGA-A4-A57E | | TCGA-AX-A0J0 | |
| TCGA-B9-4115 | | TCGA-AX-A1CF | |
| TCGA-BQ-5875 | | TCGA-AX-A1CI | |
| TCGA-BQ-5877 | | TCGA-AX-A1CK | |
| TCGA-BQ-5878 | | TCGA-AX-A2H8 | |
| TCGA-BQ-5879 | | TCGA-AX-A2HA | |
| TCGA-BQ-5882 | | TCGA-AX-A2HC | |
| TCGA-BQ-5884 | | TCGA-AX-A2HD | |
| TCGA-BQ-5887 | | TCGA-BG-A2AD | |
| TCGA-BQ-5888 | | TCGA-BG-A3EW | |
| TCGA-BQ-5890 | | TCGA-BG-A3PP | |
| TCGA-BQ-5891 | | TCGA-BK-A0CB | |
| TCGA-BQ-5894 | | TCGA-BK-A13C | |
| TCGA-BQ-7044 | | TCGA-BK-A4ZD | |
| TCGA-BQ-7045 | | TCGA-DI-A2QU | |
| TCGA-BQ-7046 | | TCGA-DI-A2QY | |
| TCGA-BQ-7051 | | TCGA-E6-A1M0 | |
| TCGA-BQ-7055 | | | |
| TCGA-BQ-7059 | | | |
| TCGA-BQ-7061 | | | |
| TCGA-DZ-6132 | | | |
| TCGA-DZ-6133 | | | |
| TCGA-DZ-6134 | | | |
| TCGA-GL-6846 | | | |
| TCGA-GL-7966 | | | |

| |
|---|
| TCGA-GL-A59R |
| TCGA-GL-A9DE |
| TCGA-P4-A5E8 |
| TCGA-P4-A5ED |
| TCGA-Y8-A8RY |

Supplementary Table S1. Data information (histone modifications)

| Sample | Cell type | Accession | Reference |
|---|---|---|---|
| lung | Somatic cell | E096 | ENCODE Project Consortium. An integrated encyclopedia of DNA elements in the human genome. Nature 2012 Sep 6;489(7414):57-74. |
| A549 | Cancer cell line | E114 | |
| liver | Somatic cell | E066 | |
| HepG2 | Cancer cell line | E118 | |

Supplementary Table S1. Data information (WGBS data)

| Abbreviation | Cancer stage | Sample ID or accession | Cancer type | Data source |
|---|---|---|---|---|
| blca_t1 | iii | TCGA-DK-A1AA-01A-11D-A23D-05 | Bladder urothelial carcinoma | TCGA |
| blca_t2 | iii | TCGA-DK-A1AG-01A-11D-A23D-05 | | |
| blca_t3 | iv | TCGA-BL-A13J-01A-11D-A23D-05 | | |
| blca_t4 | iii | TCGA-BT-A2LA-01A-11D-A23D-05 | | |
| blca_t5 | iv | TCGA-H4-A2HQ-01A-11D-A23D-05 | | |
| blca_t6 | iv | TCGA-BT-A20V-01A-11D-A23D-05 | | |
| blca_n6 | normal | TCGA-BT-A20V-11A-11D-A23D-05 | | |
| brca_t1 | iia | TCGA-A2-A04X-01A-21D-A19F-05 | Breast invasive carcinoma | |
| brca_t2 | iiia | TCGA-A8-A07I-01A-11D-A19F-05 | | |
| brca_t3 | iiic | TCGA-A2-A0YG-01A-21D-A19F-05 | | |
| brca_t4 | iia | TCGA-E2-A15H-01A-11D-A19F-05 | | |
| brca_t5 | iia | TCGA-A7-A0CE-01A-11D-A148-05 | | |
| brca_n5 | normal | TCGA-A7-A0CE-11A-21D-A148-05 | | |
| coad_t1 | i | TCGA-AA-A00R-01A-01D-A22T-05 | Colon adenocarcinoma | |
| coad_t2 | iia | TCGA-AA-3518-01A-02D-1518-05 | | |
| coad_n2 | normal | TCGA-AA-3518-11A-01D-1518-05 | | |
| gbm_t1 | - | TCGA-06-0128-01A-01D-2294-05 | Glioblastoma multiforme | |
| gbm_t2 | - | TCGA-14-1454-01A-01D-2294-05 | | |
| gbm_t3 | - | TCGA-14-3477-01A-01D-2294-05 | | |
| gbm_t4 | - | TCGA-14-1401-01A-01D-2294-05 | | |
| gbm_t5 | - | TCGA-16-1460-01A-01D-2294-05 | | |
| gbm_t6 | - | TCGA-19-1788-01A-01D-2294-05 | | |
| luad_t1 | ib | TCGA-38-4630-01A-01D-2365-05 | Lung adenocarcinoma | |
| luad_t2 | ib | TCGA-67-6215-01A-11D-2365-05 | | |
| luad_t3 | iv | TCGA-78-7156-01A-11D-2365-05 | | |
| luad_t4 | ia | TCGA-91-6840-01A-11D-2365-05 | | |

| | | | | |
|---|---|---|---|---|
| luad_t5 | ia | TCGA-44-6148-01A-11D-2365-05 | | |
| luad_n5 | normal | TCGA-44-6148-11A-01D-2365-05 | | |
| lusc_t1 | ia | TCGA-34-2600-01A-01D-1871-05 | Lung squamous cell carcinoma | |
| lusc_t2 | ib | TCGA-60-2695-01A-01D-1871-05 | | |
| lusc_t3 | ib | TCGA-21-1078-01A-01D-2365-05 | | |
| lusc_t4 | iib | TCGA-60-2722-01A-01D-1871-05 | | |
| lusc_n4 | normal | TCGA-60-2722-11A-01D-1871-05 | | |
| stad_t1 | iv | TCGA-CG-5730-01A-11D-2365-05 | Stomach adenocarcinoma | |
| stad_t2 | - | TCGA-D7-6519-01A-11D-2365-05 | | |
| stad_t3 | i | TCGA-F1-6177-01A-11D-2365-05 | | |
| stad_t4 | iia | TCGA-BR-6452-01A-12D-2365-05 | | |
| stad_n4 | normal | TCGA-BR-6452-11A-01D-2365-05 | | |
| read_t1 | iia | TCGA-AG-3593-01A-01D-2294-05 | Rectum adenocarcinoma | |
| read_t2 | iv | TCGA-AF-2689-01A-01D-2294-05 | | |
| read_n2 | normal | TCGA-AF-2689-11A-01D-2294-05 | | |
| ucec_t1 | - | TCGA-B5-A0K6-01A-11D-A23D-05 | Uterine corpus endometrial carcinoma | |
| ucec_t2 | - | TCGA-AX-A1CK-01A-11D-A23D-05 | | |
| ucec_t3 | - | TCGA-AP-A05J-01A-11D-A23D-05 | | |
| ucec_t4 | - | TCGA-A5-A0G2-01A-11D-A23D-05 | | |
| ucec_t5 | - | TCGA-AX-A1CI-01A-11D-A17H-05 | | |
| ucec_n5 | normal | TCGA-AX-A1CI-11A-11D-A17H-05 | | |
| liver_t1 | - | GSE70090 | Liver tumor and adjacent normal tissue | GEO |
| liver_t2 | - | | | |
| liver_t3 | - | | | |
| liver_t4 | - | | | |
| liver_n1 | normal | | | |
| liver_n2 | normal | | | |
| liver_n3 | normal | | | |
| liver_n4 | normal | | | |
| lung_t1 | - | | Lung tumor and adjacent normal tissue | |
| lung_t2 | - | | | |
| lung_t3 | - | | | |
| lung_n1 | normal | | | |
| lung_n2 | normal | | | |
| lung_n3 | normal | | | |
| colon_t1 | - | GSE46644 | colon moderately differentiated adenocarcinoma and adjacent normal tissue | |
| colon_n1 | normal | | | |
| colon_t2 | - | GSE52271 | Colorectal cancer and adjacent normal tissue | |
| colon_n2 | normal | | | |

| | | | |
|---|---|---|---|
| lung | normal | GSM983647 | lung cancer |
| A549 | cancer cell line | GSE127301 | |
| liver | normal | GSM916049 | liver cancer |
| HepG2 | cancer cell line | GSE46644 | |

Supplementary Table S1. Data information (Hi-C)

| Sample | Cell type | Accession | Reference |
|---|---|---|---|
| lung | Somatic cell | GSE87112 | Schmitt AD, Hu M, Jung I, Xu Z et al. A Compendium of Chromatin Contact Maps Reveals Spatially Active Regions in the Human Genome. Cell Rep.2016;17:2042-59. |
| pancreas | | | |
| H1 | Embryonic stem cell | | |
| A549 | Cancer cell line | ENCSR444WCZ | ENCODE Project Consortium. An integrated encyclopedia of DNA elements in the human genome. Nature 2012 Sep 6;489(7414):57-74. |
| Panc1 | Cancer cell line | ENCSR440CTR | |
| growing | Growing cell | PRJEB8073 | Chandra T, Ewels PA, Schoenfelder S, Furlan-Magaril M, Wingett SW, Kirschner K, Thuret JY, Andrews S, Fraser P and Reik W. Global reorganization of the nuclear landscape in senescent cells. Cell Rep. 2015;10: 471-83. |
| senescence | Senescent cell | | |
| G1 | G1 stage of mouse embryonic stem cell | GSE94489 | T. Nagano et al., Cell-cycle dynamics of chromosomal organization at single-cell resolution. Nature 547, 61 (2017). |
| early S | Early S stage of mouse embryonic stem cell | | |
