## Supplemental Table 2-13 for "Domain segregated 3D chromatin structure and segmented DNA methylation in carcinogenesis"

### Supplemental Tables

Supplementary Table S2. Amount and proportion of hypermethylated and hypomethylated CGIs.

| Sample | Stage | CGI amount | Number of hypomethylated CGIs | Proportion of hypomethylated CGIs | Number of hypermethylated CGIs (hCGIs) | Proportion of hCGIs | F-CGI amount | Number of F-hCGI | Proportion of F-hCGI | P-CGI amount | Number of P-hCGI | Proportion of P-hCGI |
|---|---|---|---|---|---|---|---|---|---|---|---|---|
| luad5 | ia | 25630 | 133 | 0.52% | 141 | 0.55% | 23635 | 118 | 0.50% | 1995 | 23 | 1.15% |
| stad4 | iia | 25646 | 389 | 1.52% | 2249 | 8.77% | 23653 | 1869 | 7.90% | 1993 | 380 | 19.07% |
| brca5 | iia | 25795 | 385 | 1.49% | 1725 | 6.69% | 23798 | 1470 | 6.18% | 1997 | 255 | 12.77% |
| coad2 | iia | 25861 | 167 | 0.65% | 4637 | 17.93% | 23858 | 3953 | 16.57% | 2003 | 684 | 34.15% |
| lusc4 | iib | 25691 | 421 | 1.64% | 4217 | 16.41% | 23706 | 3772 | 15.91% | 1985 | 445 | 22.42% |
| blca6 | iv | 25110 | 989 | 3.94% | 2454 | 9.77% | 23118 | 2145 | 9.28% | 1992 | 309 | 15.51% |
| read2 | iv | 25735 | 878 | 3.41% | 3007 | 11.68% | 23735 | 2494 | 10.51% | 2000 | 513 | 25.65% |
| liver3 | - | 24949 | 38 | 0.15% | 330 | 1.32% | 23010 | 286 | 1.24% | 1939 | 44 | 2.27% |
| liver4 | - | 24911 | 1724 | 6.92% | 1732 | 6.95% | 22977 | 1583 | 6.89% | 1934 | 149 | 7.70% |
| liver1 | - | 24926 | 56 | 0.22% | 372 | 1.49% | 22994 | 324 | 1.41% | 1932 | 48 | 2.48% |
| liver2 | - | 24968 | 2455 | 9.83% | 665 | 2.66% | 23027 | 628 | 2.73% | 1941 | 37 | 1.91% |
| colon1 | - | 26243 | 258 | 0.98% | 914 | 3.48% | 24197 | 774 | 3.20% | 2046 | 140 | 6.84% |
| colon2 | - | 25979 | 88 | 0.34% | 686 | 2.64% | 23963 | 552 | 2.30% | 2016 | 134 | 6.65% |
| lung1 | - | 25035 | 265 | 1.06% | 135 | 0.54% | 23090 | 123 | 0.53% | 1945 | 12 | 0.62% |
| lung2 | - | 24932 | 28 | 0.11% | 64 | 0.26% | 22995 | 62 | 0.27% | 1937 | 2 | 0.10% |
| lung3 | - | 25007 | 596 | 2.38% | 216 | 0.86% | 23067 | 199 | 0.86% | 1940 | 17 | 0.88% |
| ucec5 | - | 25797 | 615 | 2.38% | 3063 | 11.87% | 23796 | 2674 | 11.24% | 2001 | 389 | 19.44% |
| average | | | | 2.21% | | 6.11% | | | 5.74% | | | 10.57% |
| 103_26yr (T cell) | | 24633 | 69 | 0.28% | 238 | 0.97% | 22718 | 228 | 1.00% | 1915 | 10 | 0.52% |

Supplementary Table S3. Average CpG density, GC content, CGI length for methylation unchanged CGIs and hypomethylated CGIs.

| Sample | CpG density | | | GC content | | | CGI length | | |
|---|---|---|---|---|---|---|---|---|---|
| | unchanged CGI | hypomethylated CGI | *P*-value | unchanged CGI | hypomethylated CGI | *P*-value | unchanged CGI | hypomethylated CGI | *P*-value |
| luad5 | 0.098 | 0.089 | 1.24E-07 | 0.687 | 0.687 | 9.33E-01 | 770.5 | 350.8 | 7.47E-53 |
| stad4 | 0.098 | 0.09 | 1.13E-17 | 0.686 | 0.685 | 6.10E-01 | 761.8 | 444.7 | 4.42E-53 |
| brca5 | 0.098 | 0.086 | 3.13E-46 | 0.688 | 0.673 | 3.14E-10 | 772.1 | 403.5 | 1.55E-71 |
| coad2 | 0.097 | 0.086 | 1.74E-19 | 0.686 | 0.674 | 4.75E-04 | 722.6 | 373 | 5.29E-44 |
| lusc4 | 0.099 | 0.088 | 1.11E-26 | 0.688 | 0.68 | 1.97E-03 | 765.4 | 608 | 2.03E-04 |
| blca6 | 0.099 | 0.083 | 1.96E-210 | 0.689 | 0.664 | 1.74E-55 | 779.3 | 406.3 | 7.10E-169 |
| read2 | 0.098 | 0.082 | 1.34E-197 | 0.687 | 0.672 | 2.18E-25 | 754.4 | 357.3 | 4.17E-239 |
| liver1 | 0.098 | 0.086 | 9.51E-08 | 0.687 | 0.671 | 1.01E-02 | 773.8 | 450.1 | 7.94E-05 |
| liver3 | 0.098 | 0.086 | 2.45E-05 | 0.687 | 0.671 | 6.65E-02 | 773.5 | 381.7 | 2.38E-11 |
| liver2 | 0.099 | 0.085 | 0.00E+00 | 0.689 | 0.664 | 1.73E-148 | 807.5 | 451.3 | 1.24E-141 |
| liver4 | 0.099 | 0.084 | 3.74E-293 | 0.688 | 0.661 | 1.42E-117 | 780.5 | 417.1 | 3.74E-115 |
| lung1 | 0.098 | 0.086 | 9.03E-31 | 0.687 | 0.678 | 1.71E-03 | 772.8 | 483.3 | 3.73E-22 |
| lung3 | 0.098 | 0.084 | 7.61E-104 | 0.688 | 0.666 | 3.45E-29 | 779.6 | 386.2 | 1.85E-140 |
| lung2 | 0.098 | 0.087 | 3.41E-03 | 0.687 | 0.66 | 2.36E-03 | 768.7 | 403.6 | 6.24E-09 |
| colon1 | 0.098 | 0.088 | 7.42E-17 | 0.687 | 0.683 | 2.56E-01 | 760.3 | 413.5 | 7.17E-54 |
| colon2 | 0.098 | 0.085 | 4.15E-12 | 0.687 | 0.672 | 1.47E-03 | 762.6 | 331.4 | 1.87E-45 |
| ucec5 | 0.099 | 0.083 | 1.84E-121 | 0.688 | 0.67 | 8.66E-25 | 774.3 | 393.7 | 2.07E-149 |
| A549 | 0.098 | 0.087 | 2.46E-61 | 0.687 | 0.665 | 1.81E-29 | 754.8 | 530.9 | 1.35E-22 |
| HepG2 | 0.099 | 0.081 | 0.00E+00 | 0.688 | 0.666 | 3.31E-68 | 774.1 | 422.6 | 4.76E-25 |

Supplementary Table S4. Average brain specificity for uCGIs and hCGIs.

| Sample | Average | | *P*-value | Average | | *P*-value |
|---|---|---|---|---|---|---|
| | F-uCGI | F-hCGI | | P-uCGI | P-hCGI | |
| luad5 | 0.2411 | 0.3321 | 1.20E-01 | 0.3946 | 0.5815 | 3.96E-01 |
| stad4 | 0.2013 | 0.5957 | 4.45E-32 | 0.2668 | 0.9016 | 3.16E-16 |
| brca5 | 0.2098 | 0.6758 | 9.39E-20 | 0.3293 | 1.0433 | 8.81E-09 |
| coad2 | 0.1448 | 0.683 | 4.87E-77 | 0.2005 | 0.8119 | 3.61E-26 |
| lusc4 | 0.1774 | 0.5759 | 7.35E-47 | 0.2782 | 0.9919 | 1.53E-13 |
| blca6 | 0.1924 | 0.5747 | 4.70E-28 | 0.2869 | 1.0908 | 6.43E-13 |
| read2 | 0.1686 | 0.7678 | 5.32E-50 | 0.2119 | 0.9918 | 3.36E-27 |
| liver1 | 0.238 | 0.385 | 6.28E-03 | 0.3912 | 0.761 | 2.63E-02 |
| liver3 | 0.2353 | 0.5196 | 2.93E-04 | 0.3864 | 1.0929 | 3.19E-04 |
| liver2 | 0.2134 | 0.3204 | 3.18E-04 | 0.3774 | 0.3993 | 8.72E-01 |
| liver4 | 0.1889 | 0.6244 | 2.67E-28 | 0.361 | 0.6208 | 3.94E-03 |
| lung1 | 0.2373 | 0.2991 | 5.27E-01 | 0.3981 | 0.5386 | 7.96E-01 |
| lung3 | 0.2363 | 0.2985 | 4.89E-01 | 0.3912 | 0.7605 | 2.89E-01 |
| lung2 | 0.2393 | 0.4423 | 5.69E-02 | 0.3981 | - | - |
| colon1 | 0.2229 | 0.6431 | 2.27E-12 | 0.3636 | 0.8163 | 2.69E-06 |
| colon2 | 0.2314 | 0.5597 | 9.81E-09 | 0.3519 | 0.9204 | 5.79E-07 |
| ucec5 | 0.1906 | 0.6173 | 1.28E-31 | 0.2531 | 1.1481 | 1.62E-16 |
| A549 | 0.1703 | 0.5497 | 1.13E-47 | 0.2859 | 1.0402 | 2.23E-13 |
| HepG2 | 0.1874 | 0.4916 | 4.84E-29 | 0.3179 | 0.7555 | 2.44E-08 |

Supplementary Table S5. The proportion of compartment B in normal and tumorous samples (chr1 to chr22).

| | Normal sample | Tumorous cell line |
|---|---|---|
| Pancreas | 0.5222 | 0.6205 |
| Lung | 0.5548 | 0.5718 |

Supplementary Table S6. Averaged compartment vector $\bar{V}$ in regions with different sequential properties (chr1).

| | | | Normal sample | Tumorous cell line | Difference[a] |
|---|---|---|---|---|---|
| Lung | forest | CGI | 0.0205 | 0.0159 | -0.0046 |
| | | nonCGI | 0.0089 | 0.0058 | -0.0031 |
| | | Difference[b] | 0.0115 | 0.01 | - |
| | prairie | CGI | -0.0183 | -0.0256 | -0.0073 |
| | | nonCGI | -0.0247 | -0.0269 | -0.0022 |
| | | Difference | 0.0064 | 0.0013 | - |
| Pancreas | forest | CGI | 0.0236 | 0.0126 | -0.011 |
| | | nonCGI | 0.013 | 0.001 | -0.012 |

| | | | | | |
|---|---|---|---|---|---|
| | | Difference | 0.0106 | 0.0116 | - |
| | prairie | CGI | -0.0091 | -0.0308 | -0.0218 |
| | | nonCGI | -0.0133 | -0.0283 | -0.015 |
| | | Difference | 0.0042 | -0.0025 | - |

a. The sample difference is defined as the averaged compartment vector $\bar{V}$ of tumorous cell line minus that of normal sample in the same region.

b. The region difference is defined as the $\bar{V}$ of CGI regions minus that of nonCGI regions, and is calculated for forest and prairie respectively.

Supplementary Table S7. GO enrichment analysis of aggregated F-CGI genes with increased gene expression in A549.

| ID | Description | p.adjust |
|---|---|---|
| GO:0034660 | ncRNA metabolic process | 1.28E-12 |
| GO:0006415 | translational termination | 1.71E-10 |
| GO:0043624 | cellular protein complex disassembly | 5.94E-08 |
| GO:0140053 | mitochondrial gene expression | 6.07E-08 |
| GO:0042254 | ribosome biogenesis | 2.10E-05 |
| GO:0006353 | DNA-templated transcription, termination | 2.18E-05 |
| GO:0019080 | viral gene expression | 5.70E-05 |
| GO:0034404 | nucleobase-containing small molecule biosynthetic process | 8.19E-05 |
| GO:0032200 | telomere organization | 1.81E-04 |
| GO:0000723 | telomere maintenance | 1.81E-04 |
| GO:0007059 | chromosome segregation | 2.33E-04 |
| GO:0051169 | nuclear transport | 2.71E-04 |
| GO:0000280 | nuclear division | 3.89E-04 |
| GO:1902749 | regulation of cell cycle G2/M phase transition | 1.08E-03 |
| GO:0048193 | Golgi vesicle transport | 1.24E-03 |
| GO:0044839 | cell cycle G2/M phase transition | 1.41E-03 |
| GO:0072594 | establishment of protein localization to organelle | 1.41E-03 |
| GO:0140014 | mitotic nuclear division | 3.97E-03 |
| GO:0005996 | monosaccharide metabolic process | 4.29E-03 |
| GO:0006401 | RNA catabolic process | 4.37E-03 |
| GO:0016925 | protein sumoylation | 4.37E-03 |
| GO:0046434 | organophosphate catabolic process | 4.55E-03 |
| GO:0071824 | protein-DNA complex subunit organization | 5.13E-03 |
| GO:0006096 | glycolytic process | 5.93E-03 |
| GO:0006403 | RNA localization | 6.24E-03 |
| GO:0034504 | protein localization to nucleus | 6.53E-03 |
| GO:0009100 | glycoprotein metabolic process | 6.65E-03 |
| GO:0033044 | regulation of chromosome organization | 8.08E-03 |
| GO:0002566 | somatic diversification of immune receptors via somatic mutation | 8.26E-03 |
| GO:0042866 | pyruvate biosynthetic process | 9.62E-03 |

| | | |
|---|---|---|
| GO:0006457 | protein folding | 1.14E-02 |
| GO:0019320 | hexose catabolic process | 1.15E-02 |
| GO:0051310 | metaphase plate congression | 1.15E-02 |
| GO:0097064 | ncRNA export from nucleus | 1.21E-02 |
| GO:0050000 | chromosome localization | 1.21E-02 |
| GO:0000070 | mitotic sister chromatid segregation | 1.59E-02 |
| GO:0065004 | protein-DNA complex assembly | 1.70E-02 |
| GO:0015931 | nucleobase-containing compound transport | 1.82E-02 |
| GO:1900034 | regulation of cellular response to heat | 1.86E-02 |
| GO:0006733 | oxidoreduction coenzyme metabolic process | 1.92E-02 |
| GO:0006611 | protein export from nucleus | 1.97E-02 |
| GO:0006486 | protein glycosylation | 2.05E-02 |
| GO:0051321 | meiotic cell cycle | 2.05E-02 |
| GO:0046390 | ribose phosphate biosynthetic process | 2.05E-02 |
| GO:0000076 | DNA replication checkpoint | 2.05E-02 |
| GO:0097711 | ciliary basal body-plasma membrane docking | 2.58E-02 |
| GO:0006090 | pyruvate metabolic process | 2.63E-02 |
| GO:0044766 | multi-organism transport | 2.63E-02 |
| GO:1902579 | multi-organism localization | 2.63E-02 |
| GO:0000226 | microtubule cytoskeleton organization | 3.17E-02 |
| GO:0060968 | regulation of gene silencing | 3.19E-02 |
| GO:0075733 | intracellular transport of virus | 3.34E-02 |
| GO:0072524 | pyridine-containing compound metabolic process | 3.38E-02 |
| GO:0016052 | carbohydrate catabolic process | 3.56E-02 |
| GO:0006901 | vesicle coating | 4.05E-02 |
| GO:0045815 | positive regulation of gene expression, epigenetic | 5.38E-02 |
| GO:0060122 | inner ear receptor cell stereocilium organization | 5.38E-02 |
| GO:0008089 | anterograde axonal transport | 5.44E-02 |
| GO:0034728 | nucleosome organization | 5.81E-02 |
| GO:0048208 | COPII vesicle coating | 5.92E-02 |
| GO:1900182 | positive regulation of protein localization to nucleus | 5.92E-02 |
| GO:0071166 | ribonucleoprotein complex localization | 6.46E-02 |
| GO:0043470 | regulation of carbohydrate catabolic process | 6.56E-02 |
| GO:0006293 | nucleotide-excision repair, preincision complex stabilization | 8.04E-02 |
| GO:0043467 | regulation of generation of precursor metabolites and energy | 8.36E-02 |
| GO:0019692 | deoxyribose phosphate metabolic process | 8.79E-02 |

Supplementary Table S8. GO enrichment analysis of aggregated P-CGI genes with increased gene expression in A549.

| ID | Description | p.adjust |
|---|---|---|
| GO:0007492 | endoderm development | 2.69E-02 |
| GO:0007223 | Wnt signaling pathway, calcium modulating pathway | 2.69E-02 |
| GO:0030488 | tRNA methylation | 2.69E-02 |

| | | |
|---|---|---|
| GO:0000280 | nuclear division | 2.69E-02 |
| GO:0072498 | embryonic skeletal joint development | 2.95E-02 |
| GO:0001711 | endodermal cell fate commitment | 3.13E-02 |
| GO:0048641 | regulation of skeletal muscle tissue development | 3.13E-02 |
| GO:0007059 | chromosome segregation | 3.13E-02 |
| GO:0140053 | mitochondrial gene expression | 3.13E-02 |
| GO:0030900 | forebrain development | 4.45E-02 |
| GO:0140014 | mitotic nuclear division | 5.56E-02 |
| GO:0009163 | nucleoside biosynthetic process | 6.85E-02 |
| GO:1901659 | glycosyl compound biosynthetic process | 6.96E-02 |
| GO:0046677 | response to antibiotic | 7.10E-02 |
| GO:0060272 | embryonic skeletal joint morphogenesis | 7.10E-02 |
| GO:0030326 | embryonic limb morphogenesis | 7.10E-02 |
| GO:0035113 | embryonic appendage morphogenesis | 7.10E-02 |
| GO:0060394 | negative regulation of pathway-restricted SMAD protein phosphorylation | 7.89E-02 |
| GO:0000075 | cell cycle checkpoint | 9.16E-02 |
| GO:0070498 | interleukin-1-mediated signaling pathway | 9.16E-02 |
| GO:2000772 | regulation of cellular senescence | 9.24E-02 |
| GO:0090307 | mitotic spindle assembly | 9.24E-02 |

Supplementary Table S9. GO enrichment analysis of aggregated F-CGI genes with increased gene expression in Panc1.

| ID | Description | p.adjust |
|---|---|---|
| GO:0048193 | Golgi vesicle transport | 3.56E-04 |
| GO:0007156 | homophilic cell adhesion via plasma membrane adhesion molecules | 3.56E-04 |
| GO:0048562 | embryonic organ morphogenesis | 4.95E-04 |
| GO:0048568 | embryonic organ development | 4.95E-04 |
| GO:0003002 | regionalization | 1.90E-03 |
| GO:0006643 | membrane lipid metabolic process | 1.90E-03 |
| GO:0006506 | GPI anchor biosynthetic process | 3.62E-03 |
| GO:0022411 | cellular component disassembly | 7.03E-03 |
| GO:0000226 | microtubule cytoskeleton organization | 1.22E-02 |
| GO:0051656 | establishment of organelle localization | 1.22E-02 |
| GO:0007163 | establishment or maintenance of cell polarity | 1.22E-02 |
| GO:0140014 | mitotic nuclear division | 1.50E-02 |
| GO:0006735 | NADH regeneration | 1.50E-02 |
| GO:0061621 | canonical glycolysis | 1.50E-02 |
| GO:0061718 | glucose catabolic process to pyruvate | 1.50E-02 |
| GO:0048285 | organelle fission | 1.56E-02 |
| GO:0050804 | modulation of chemical synaptic transmission | 1.89E-02 |
| GO:0099177 | regulation of trans-synaptic signaling | 2.00E-02 |

| | | |
|---|---|---|
| GO:1905477 | positive regulation of protein localization to membrane | 2.20E-02 |
| GO:1990778 | protein localization to cell periphery | 2.64E-02 |
| GO:0048667 | cell morphogenesis involved in neuron differentiation | 2.71E-02 |
| GO:0008589 | regulation of smoothened signaling pathway | 3.83E-02 |
| GO:0010975 | regulation of neuron projection development | 3.99E-02 |
| GO:0021953 | central nervous system neuron differentiation | 4.03E-02 |
| GO:0007018 | microtubule-based movement | 4.06E-02 |
| GO:0010769 | regulation of cell morphogenesis involved in differentiation | 4.06E-02 |
| GO:0043254 | regulation of protein complex assembly | 4.13E-02 |
| GO:0072659 | protein localization to plasma membrane | 4.13E-02 |
| GO:2001233 | regulation of apoptotic signaling pathway | 4.13E-02 |
| GO:0045787 | positive regulation of cell cycle | 4.44E-02 |
| GO:0018212 | peptidyl-tyrosine modification | 4.59E-02 |
| GO:0032506 | cytokinetic process | 4.88E-02 |
| GO:0010498 | proteasomal protein catabolic process | 4.93E-02 |
| GO:1905475 | regulation of protein localization to membrane | 4.98E-02 |
| GO:0019886 | antigen processing and presentation of exogenous peptide antigen via MHC class II | 5.02E-02 |
| GO:0070085 | glycosylation | 5.20E-02 |
| GO:0006020 | inositol metabolic process | 5.26E-02 |
| GO:0008608 | attachment of spindle microtubules to kinetochore | 5.34E-02 |
| GO:0032350 | regulation of hormone metabolic process | 5.39E-02 |
| GO:0051591 | response to cAMP | 5.39E-02 |
| GO:0035418 | protein localization to synapse | 5.39E-02 |
| GO:0007416 | synapse assembly | 5.63E-02 |
| GO:0021700 | developmental maturation | 5.69E-02 |
| GO:0023061 | signal release | 6.03E-02 |
| GO:0016052 | carbohydrate catabolic process | 6.03E-02 |
| GO:0060562 | epithelial tube morphogenesis | 6.04E-02 |
| GO:0099504 | synaptic vesicle cycle | 6.04E-02 |
| GO:0006415 | translational termination | 6.04E-02 |
| GO:0046165 | alcohol biosynthetic process | 6.04E-02 |
| GO:0099003 | vesicle-mediated transport in synapse | 6.04E-02 |
| GO:0032355 | response to estradiol | 6.04E-02 |
| GO:0045185 | maintenance of protein location | 6.60E-02 |
| GO:0006334 | nucleosome assembly | 6.64E-02 |
| GO:0071103 | DNA conformation change | 6.77E-02 |
| GO:0006457 | protein folding | 6.77E-02 |
| GO:1905383 | protein localization to presynapse | 6.77E-02 |
| GO:0030866 | cortical actin cytoskeleton organization | 6.77E-02 |
| GO:1902186 | regulation of viral release from host cell | 6.77E-02 |
| GO:0006914 | autophagy | 6.77E-02 |
| GO:0061919 | process utilizing autophagic mechanism | 6.77E-02 |

| | | |
|---|---|---|
| GO:0034728 | nucleosome organization | 6.83E-02 |
| GO:0009100 | glycoprotein metabolic process | 7.05E-02 |
| GO:0031667 | response to nutrient levels | 7.15E-02 |
| GO:0099111 | microtubule-based transport | 7.34E-02 |
| GO:0051648 | vesicle localization | 7.49E-02 |
| GO:0000070 | mitotic sister chromatid segregation | 7.53E-02 |
| GO:0006575 | cellular modified amino acid metabolic process | 7.54E-02 |
| GO:0007088 | regulation of mitotic nuclear division | 7.67E-02 |
| GO:0006497 | protein lipidation | 7.67E-02 |
| GO:0008306 | associative learning | 7.67E-02 |
| GO:0006486 | protein glycosylation | 7.67E-02 |
| GO:0048265 | response to pain | 8.25E-02 |
| GO:0099173 | postsynapse organization | 8.75E-02 |
| GO:0009812 | flavonoid metabolic process | 8.75E-02 |
| GO:0045472 | response to ether | 8.75E-02 |
| GO:0051988 | regulation of attachment of spindle microtubules to kinetochore | 8.75E-02 |
| GO:0070482 | response to oxygen levels | 8.79E-02 |
| GO:0006590 | thyroid hormone generation | 9.36E-02 |
| GO:0042073 | intraciliary transport | 9.63E-02 |
| GO:0106027 | neuron projection organization | 9.81E-02 |
| GO:0070723 | response to cholesterol | 9.81E-02 |
| GO:0031345 | negative regulation of cell projection organization | 9.92E-02 |
| GO:0032535 | regulation of cellular component size | 9.97E-02 |

Supplementary Table S10. GO enrichment analysis of aggregated P-CGI genes with increased gene expression in Panc1.

| ID | Description | p.adjust |
|---|---|---|
| GO:0010717 | regulation of epithelial to mesenchymal transition | 3.04E-05 |
| GO:0035107 | appendage morphogenesis | 3.04E-05 |
| GO:0035108 | limb morphogenesis | 3.04E-05 |
| GO:0030326 | embryonic limb morphogenesis | 3.04E-05 |
| GO:0035113 | embryonic appendage morphogenesis | 3.04E-05 |
| GO:0007492 | endoderm development | 1.24E-04 |
| GO:2000826 | regulation of heart morphogenesis | 1.57E-04 |
| GO:0048762 | mesenchymal cell differentiation | 1.89E-03 |
| GO:0001837 | epithelial to mesenchymal transition | 2.29E-03 |
| GO:0016202 | regulation of striated muscle tissue development | 2.88E-03 |
| GO:0010463 | mesenchymal cell proliferation | 2.90E-03 |
| GO:1901861 | regulation of muscle tissue development | 2.92E-03 |
| GO:1904837 | beta-catenin-TCF complex assembly | 3.66E-03 |
| GO:0061311 | cell surface receptor signaling pathway involved in heart development | 4.04E-03 |
| GO:0035690 | cellular response to drug | 5.49E-03 |

| | | |
|---|---|---|
| GO:0110110 | positive regulation of animal organ morphogenesis | 6.36E-03 |
| GO:0009116 | nucleoside metabolic process | 7.98E-03 |
| GO:0010560 | positive regulation of glycoprotein biosynthetic process | 7.98E-03 |
| GO:0001649 | osteoblast differentiation | 8.29E-03 |
| GO:0001503 | ossification | 8.29E-03 |
| GO:0045667 | regulation of osteoblast differentiation | 1.30E-02 |
| GO:0030278 | regulation of ossification | 1.30E-02 |
| GO:1901657 | glycosyl compound metabolic process | 1.42E-02 |
| GO:0090090 | negative regulation of canonical Wnt signaling pathway | 1.45E-02 |
| GO:0030201 | heparan sulfate proteoglycan metabolic process | 1.45E-02 |
| GO:0010038 | response to metal ion | 1.49E-02 |
| GO:0010001 | glial cell differentiation | 1.58E-02 |
| GO:0072078 | nephron tubule morphogenesis | 1.58E-02 |
| GO:0060394 | negative regulation of pathway-restricted SMAD protein phosphorylation | 1.66E-02 |
| GO:0050808 | synapse organization | 1.80E-02 |
| GO:0198738 | cell-cell signaling by wnt | 2.11E-02 |
| GO:0007548 | sex differentiation | 2.15E-02 |
| GO:0070542 | response to fatty acid | 2.37E-02 |
| GO:0007568 | aging | 2.37E-02 |
| GO:0031128 | developmental induction | 2.44E-02 |
| GO:0010464 | regulation of mesenchymal cell proliferation | 2.63E-02 |
| GO:0050679 | positive regulation of epithelial cell proliferation | 2.69E-02 |
| GO:0098727 | maintenance of cell number | 2.69E-02 |
| GO:0071634 | regulation of transforming growth factor beta production | 2.91E-02 |
| GO:0016572 | histone phosphorylation | 3.21E-02 |
| GO:0042149 | cellular response to glucose starvation | 3.21E-02 |
| GO:0071604 | transforming growth factor beta production | 3.21E-02 |
| GO:0006687 | glycosphingolipid metabolic process | 3.30E-02 |
| GO:1904888 | cranial skeletal system development | 3.30E-02 |
| GO:1901652 | response to peptide | 3.34E-02 |
| GO:0032288 | myelin assembly | 3.34E-02 |
| GO:0034063 | stress granule assembly | 3.34E-02 |
| GO:0050804 | modulation of chemical synaptic transmission | 3.38E-02 |
| GO:0099177 | regulation of trans-synaptic signaling | 3.41E-02 |
| GO:0060973 | cell migration involved in heart development | 3.63E-02 |
| GO:0031641 | regulation of myelination | 3.73E-02 |
| GO:1901659 | glycosyl compound biosynthetic process | 3.92E-02 |
| GO:0043087 | regulation of GTPase activity | 4.10E-02 |
| GO:0036499 | PERK-mediated unfolded protein response | 4.25E-02 |
| GO:1901522 | positive regulation of transcription from RNA polymerase II promoter involved in cellular response to chemical stimulus | 4.53E-02 |
| GO:0097305 | response to alcohol | 4.70E-02 |
| GO:0046677 | response to antibiotic | 5.03E-02 |

| | | |
|---|---|---|
| GO:0001570 | vasculogenesis | 5.10E-02 |
| GO:0002762 | negative regulation of myeloid leukocyte differentiation | 5.13E-02 |
| GO:0042180 | cellular ketone metabolic process | 5.43E-02 |
| GO:0043547 | positive regulation of GTPase activity | 5.75E-02 |
| GO:0022407 | regulation of cell-cell adhesion | 5.75E-02 |
| GO:0072528 | pyrimidine-containing compound biosynthetic process | 5.99E-02 |
| GO:0046661 | male sex differentiation | 5.99E-02 |
| GO:0031645 | negative regulation of neurological system process | 5.99E-02 |
| GO:2000679 | positive regulation of transcription regulatory region DNA binding | 5.99E-02 |
| GO:0019229 | regulation of vasoconstriction | 6.28E-02 |
| GO:0042304 | regulation of fatty acid biosynthetic process | 6.55E-02 |
| GO:0042326 | negative regulation of phosphorylation | 6.89E-02 |
| GO:0097306 | cellular response to alcohol | 6.89E-02 |
| GO:0051668 | localization within membrane | 6.96E-02 |
| GO:0071248 | cellular response to metal ion | 6.97E-02 |
| GO:0071398 | cellular response to fatty acid | 7.09E-02 |
| GO:0071333 | cellular response to glucose stimulus | 7.24E-02 |
| GO:0006929 | substrate-dependent cell migration | 7.24E-02 |
| GO:0035418 | protein localization to synapse | 7.24E-02 |
| GO:0051055 | negative regulation of lipid biosynthetic process | 7.24E-02 |
| GO:1901655 | cellular response to ketone | 7.26E-02 |
| GO:0031346 | positive regulation of cell projection organization | 7.59E-02 |
| GO:0051194 | positive regulation of cofactor metabolic process | 7.60E-02 |
| GO:0060325 | face morphogenesis | 7.60E-02 |
| GO:0001840 | neural plate development | 7.68E-02 |
| GO:0006930 | substrate-dependent cell migration, cell extension | 7.68E-02 |
| GO:0060287 | epithelial cilium movement involved in determination of left/right asymmetry | 7.68E-02 |
| GO:0070601 | centromeric sister chromatid cohesion | 7.68E-02 |
| GO:0097050 | type B pancreatic cell apoptotic process | 7.68E-02 |
| GO:0106049 | regulation of cellular response to osmotic stress | 7.68E-02 |
| GO:2000665 | regulation of interleukin-13 secretion | 7.68E-02 |
| GO:2000767 | positive regulation of cytoplasmic translation | 7.68E-02 |
| GO:2001225 | regulation of chloride transport | 7.68E-02 |
| GO:0033687 | osteoblast proliferation | 7.78E-02 |
| GO:0072527 | pyrimidine-containing compound metabolic process | 7.84E-02 |
| GO:0046434 | organophosphate catabolic process | 8.14E-02 |
| GO:0040036 | regulation of fibroblast growth factor receptor signaling pathway | 8.14E-02 |
| GO:0050673 | epithelial cell proliferation | 8.21E-02 |
| GO:0042593 | glucose homeostasis | 8.45E-02 |
| GO:0072611 | interleukin-13 secretion | 8.45E-02 |
| GO:0090042 | tubulin deacetylation | 8.45E-02 |
| GO:1904322 | cellular response to forskolin | 8.45E-02 |

| | | |
|---|---|---|
| GO:0097009 | energy homeostasis | 9.03E-02 |
| GO:0006352 | DNA-templated transcription, initiation | 9.03E-02 |
| GO:0007050 | cell cycle arrest | 9.03E-02 |
| GO:0014823 | response to activity | 9.14E-02 |
| GO:0001678 | cellular glucose homeostasis | 9.15E-02 |
| GO:0061564 | axon development | 9.29E-02 |
| GO:0035641 | locomotory exploration behavior | 9.29E-02 |
| GO:0060788 | ectodermal placode formation | 9.29E-02 |
| GO:0072578 | neurotransmitter-gated ion channel clustering | 9.29E-02 |
| GO:0045785 | positive regulation of cell adhesion | 9.64E-02 |
| GO:0048511 | rhythmic process | 9.88E-02 |
| GO:0043647 | inositol phosphate metabolic process | 9.93E-02 |

Supplementary Table S11. The proportion of antigen processing presentation genes in F-F contact break genes. (F antigen genes take 1.26% of F genes)

| | All F antigen genes in Hi-C | F genes loss F contact at 600K-2M | Antigen genes | Proportion of FF loss genes | Proportion of antigen genes |
|---|---|---|---|---|---|
| LG2_LGt | 155 | 3329 | 46 | 1.38% | 29.68% |
| PA2_Panc1 | 157 | 3272 | 44 | 1.34% | 28.03% |
| common | | 1541 | 27 | 1.75% | |

Supplementary Table S12. The proportion of immune system genes in F-F contact break genes. (F immune genes take 13.91% of F genes)

| | All F immune genes in Hi-C | F genes loss F contact at 600K-2M | Immune system genes | Proportion of FF loss genes | Proportion of immune genes |
|---|---|---|---|---|---|
| LG2_LGt | 1893 | 3329 | 496 | 14.90% | 26.20% |
| PA2_Panc1 | 1893 | 3272 | 502 | 15.34% | 26.52% |
| common | | 1541 | 222 | 14.41% | |

Supplementary Table S13. Expression level for hCGI genes.

| Sample | Genes | Mean expression in normal | Mean expression in cancer | Mean expression changes | Number of up-regulated genes | Number of down-regulated genes |
|---|---|---|---|---|---|---|
| brca5 | F-hCGI | 7515.27 | 7230.29 | -284.98 | 321 | 407 |
| coad2 | F-hCGI | 7026.12 | 6210.48 | -815.64 | 819 | 1296 |
| luad5 | F-hCGI | 10407.17 | 10503.67 | 96.5 | 43 | 42 |
| ucec5 | F-hCGI | 7954.84 | 7526.53 | -428.31 | 610 | 717 |
| brca5 | P-hCGI | 5896.82 | 4908.3 | -988.53 | 37 | 78 |
| coad2 | P-hCGI | 6137.97 | 4419.07 | -1718.9 | 86 | 297 |
| luad5 | P-hCGI | 9718.4 | 9947.7 | 229.3 | 6 | 4 |
| ucec5 | P-hCGI | 6054.25 | 4908.42 | -1145.83 | 70 | 119 |
